# Cleavage of Braun lipoprotein Lpp from the bacterial peptidoglycan by a paralog of L,D-transpeptidases, LdtF

**DOI:** 10.1101/2021.02.24.432682

**Authors:** Raj Bahadur, Pavan Kumar Chodisetti, Manjula Reddy

## Abstract

Gram-negative bacterial cell envelope is made up of an outer membrane (OM), an inner membrane (IM) that surrounds the cytoplasm, and a periplasmic space between the two membranes containing peptidoglycan (PG or murein). PG is an elastic polymer that forms a mesh-like sacculus around the IM protecting cells from turgor and environmental stress conditions. In several bacteria including *E. coli*, the OM is tethered to PG by an abundant OM lipoprotein, Lpp (or Braun lipoprotein) that functions to maintain the structural and functional integrity of the cell envelope. Since its discovery Lpp has been studied extensively and although L,D-transpeptidases, the enzymes that catalyse the formation of PG–Lpp linkages have been earlier identified, it is not known how these linkages are modulated. Here, using genetic and biochemical approaches, we show that LdtF (formerly *yafK*), a newly-identified paralog of L,D-transpeptidases in *E. coli* is a murein hydrolytic enzyme that catalyses cleavage of Lpp from the PG sacculus. LdtF also exhibits glycine-specific carboxypeptidase activity on muropeptides containing a terminal glycine residue. LdtF is earlier presumed to be an L,D-transpeptidase; however, our results show that it is indeed an L,D-endopeptidase that hydrolyses the products generated by the L,D-transpeptidases. To summarize, this study describes the discovery of a murein endopeptidase with a hitherto unknown catalytic specificity that removes the PG–Lpp cross-links suggesting a role for LdtF in regulation of PG-OM linkages to maintain the structural integrity of the bacterial cell envelope.

**Significance statement:** Bacterial cell walls contain a unique protective exoskeleton, peptidoglycan, which is a target of several clinically important antimicrobials. In Gram-negative bacteria, peptidoglycan is covered by an additional lipid layer, outer membrane that serves as permeability barrier against entry of toxic molecules. In some bacteria, an extremely abundant lipoprotein, Lpp staples outer membrane to peptidoglycan to maintain the structural integrity of the cell envelope. In this study, we identify a previously unknown peptidoglycan hydrolytic enzyme that cleaves Lpp from the peptidoglycan sacculus and show how the outer membrane-peptidoglycan linkages are modulated in *Escherichia coli*. Overall, this study helps in understanding the fundamental bacterial cell wall biology and in identification of alternate drug targets for development of new antimicrobials.

## Introduction

Gram-negative bacterial cell envelope is made up of outer membrane (or OM), an asymmetric bilayered lipid membrane which is surface-exposed and an inner membrane (or IM) consisting of a phospholipid bilayer surrounding the cytoplasm. In between these two membranes is the periplasmic space in which a sac-like molecule, the peptidoglycan (PG or murein) is located (1). PG is an elastic heteropolymer that protects bacterial cells from lysis by internal osmotic pressure and from external stress conditions. It is a single, large macromolecule made up of multiple linear glycan strands that are inter-connected with each other by short peptide chains forming a net-like sacculus around the cytoplasmic membrane (Fig.1). The glycan strands are polymers of alternating β-1,4-linked N-acetylglucosamine (GlcNAc) and N-acetylmuramic acid (MurNAc) disaccharide units in which the D-lactoyl moiety of each MurNAc residue is covalently attached to the first amino acid of the stem peptide. Normally, the peptide chains are of two to five amino acids with a pentapeptide made up of L-alanine (L-ala^1^)–D-glutamic acid (D-glu^2^)–mesodiaminopimelic acid (mDAP^3^)–D-ala^4^–D-ala^5^ residues. In *E. coli*, approximately 40% of the neighbouring peptide chains are linked to each other, either between the D-ala^4^ and mDAP^3^ (D-ala–mDAP or 4–3) or two mDAP^3^ residues (mDAP–mDAP or 3–3) residues. Of these, the 4–3 cross-links are more prevalent and are formed by D,D-transpeptidases whereas 3–3 cross-links are much less abundant and are catalysed by L,D-transpeptidases (LDTs), LdtD and LdtE (2–4).

**Fig. 1.**
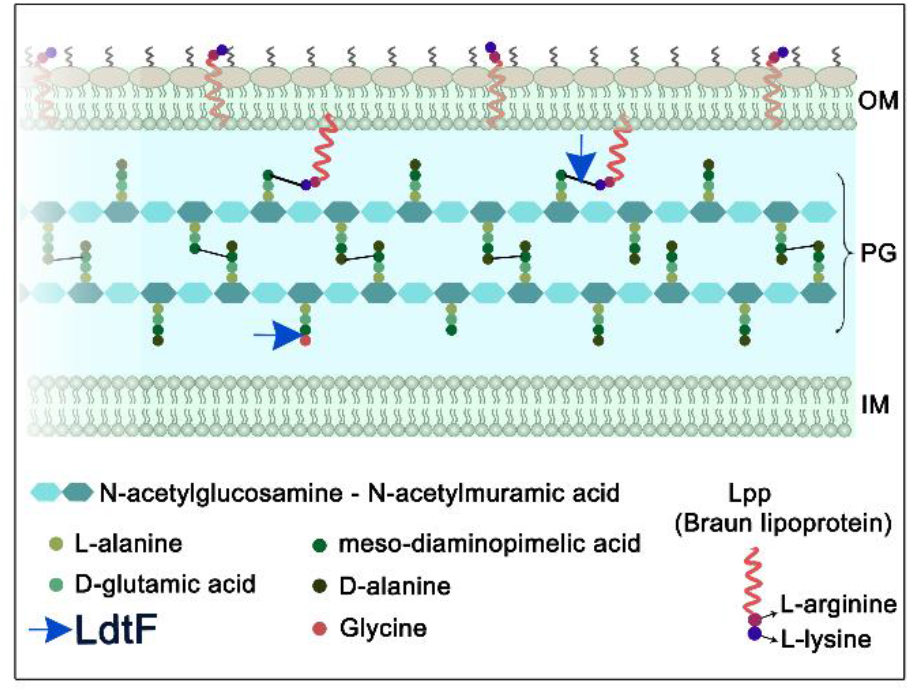
Diagrammatic representation of the cell envelope of *E. coli*. Cell envelope consists of three layers-outer membrane (OM), peptidoglycan (PG) and inner membrane (IM). PG is stapled to the OM by Lpp or Braun lipoprotein (red helix) which exists in bound or free form (5–9). In the bound form, the N-terminal end of Lpp is anchored to the OM whereas the C-terminal lysine (purple circle) is covalently attached to an mDAP residue (dark green) of the PG stem peptides. The free form of Lpp spans the OM and is exposed to the surface (11). LdtF is identified in this study as an endopeptidase which cleaves PG–Lpp cross-links and also as a glycine-specific carboxypeptidase.

In several bacteria including *E. coli*, the OM is stapled to PG by a lipoprotein, Lpp or Braun lipoprotein. Lpp is the first OM lipoprotein to be discovered almost five decades ago and has been studied extensively (5–9). It is a small abundant protein (∼10^6^ molecules per cell) of 58 amino acids and is known to exist in two conformations each occupying a distinct subcellular location in the cell envelope (9–11). One third of total Lpp is in the periplasm covalently attached to the mDAP^3^ residues of the PG peptides (bound form) whereas the other two third spans the OM (free form) (Fig.1). OM-PG tethering by Lpp has been shown to determine the width of the periplasm by controlling the IM-OM distance and to contribute to the stiffness of cell envelope (12,13). Although Lpp is not essential for viability of *E. coli*, mutants that lack Lpp show several pleiotropic defects such as increased sensitivity to hydrophobic agents, leakage of periplasmic contents, OM blebbing, vesiculation, cell separation defects, as well as deficiency in virulence, highlighting the role of Lpp in maintenance of envelope integrity (8,9,14).

Three redundant LDTs, LdtA, -B, -C catalyse the formation of PG–Lpp cross-links by covalently attaching the extreme C-terminal residue of Lpp, lysine to the mDAP^3^ residue of a tetrapeptide in the mature PG sacculus (15). In this reaction, the terminal D-ala residue of the tetrapeptide is lost leading to the formation of a tripeptide-Lpp cross-link (Fig. 1). About, 10% of the peptides in a PG sacculus are attached to Lpp and this frequency is presumed to vary during conditions of stress (2,5,16–18).

*E. coli* encodes six LDTs, LdtA-F belonging to YkuD family of proteins (3,18,19,20). Of these, LdtA,-B,-C catalyze the covalent attachment of Lpp to the PG; though these are redundant, LdtB is physiologically relevant because deleting *ldtB* alone abrogates the attachment of Lpp to a significant extent (15). Ribosome profiling (10) has also shown the abundance of LdtB to be much higher (∼5,000 copies per cell) than LdtA and LdtC (∼50 and 500 respectively). On the other hand, the 3–3 cross-link formation in the PG sacculus is catalysed by LdtD and LdtE (21).

Apart from their ability to form cross-links, LdtA-E catalyse an amino acid exchange reaction in the periplasm wherein the terminal D-ala^4^ residue of the stem peptides is substituted with either glycine or a variety of noncanonical D-amino acids (NCDAA) such as D-tryptophan, D-methionine or D-aspartate (2,21,22). The significance of this exchange in *E. coli* is not clear, although, it is believed that these substituted muropeptides do not participate in further steps of PG polymerization. LDTs are presumed to have a larger role in the maintenance of structural integrity of PG because of their ability to form cross-links de novo in a mature PG sacculus independent of active PG precursor synthesis (18).

LdtF is a recently-identified paralog of LDTs and is implicated in facilitating the formation of 3–3 cross-links; however, its precise function remains unclear (18–20). Here, we show that LdtF (encoded by *yafK*) is a murein L,D-endopeptidase that cleaves Lpp from the PG sacculus. We initially identified *ldtF* because of its genetic interaction with *mepS*, a gene encoding a major PG elongation-specific D,D-endopeptidase (23). Further genetic and biochemical analysis demonstrated the role of LdtF in hydrolysing the products generated by the activity of other LDTs. LdtF cleaves Lpp which is bound to the PG sacculus and in addition, cleaves the terminal glycine residue that is incorporated into the stem peptides due to the periplasmic exchange reaction of LDTs. However, LdtF was not able to cleave the terminal NCDAA residues from the muropeptides. To summarize, this study identifies a murein endopeptidase with a previously unknown catalytic specificity having an ability to modulate the Lpp-mediated OM-PG linkages.

## Results

### Lack of PG-Lpp linkages confer growth advantage to an *E. coli* mutant lacking an elongation-specific D,D-endopeptidase, *mepS*

We showed earlier that absence of 3–3 cross-link forming LDTs (*ldtD* and *ldtE*) confer growth advantage to a mutant lacking an elongation-specific 4–3 cross-link cleaving D,D-endopeptidase, MepS signifying the importance of cleavage of both 4–3 and 3–3 cross-links to make space for incorporation of new PG material during cell elongation (23,24). To examine whether the tethering of OM to PG by Lpp also affects the process of PG expansion, we introduced a deletion of Lpp into a mutant lacking MepS and examined the growth phenotypes of the double mutant on Nutrient agar (NA) because *mepS* deletion mutant is unable to grow on NA (25). Fig. 2A shows that absence of Lpp restores moderate amount of growth to the *mepS* deletion mutant on NA. In addition, a mutant Lpp allele that lacks the C-terminal lysine residue and hence unable to bind PG (Lpp^ΔK58^; 11) confers suppression like that of Lpp deletion. Likewise, deletion of LdtB, which catalyses the formation of mDAP–Lpp linkages conferred growth to *mepS* deletion mutant (Fig. 2A; Fig. S1, Fig. S2). However, deletion of LdtA or LdtC which also link Lpp to mDAP did not have any effect on *mepS* growth (Fig. S1). Surprisingly, deletion of LdtF, the newly-identified paralog of LDTs, conferred a very small yet consistent growth defect to *mepS* single mutant (Fig. S1) which was further exacerbated in a *mepS* mutant lacking *mepK*, a gene encoding the 3–3 cross-link cleaving PG hydrolase (Fig. 2B).

**Fig. 2.**
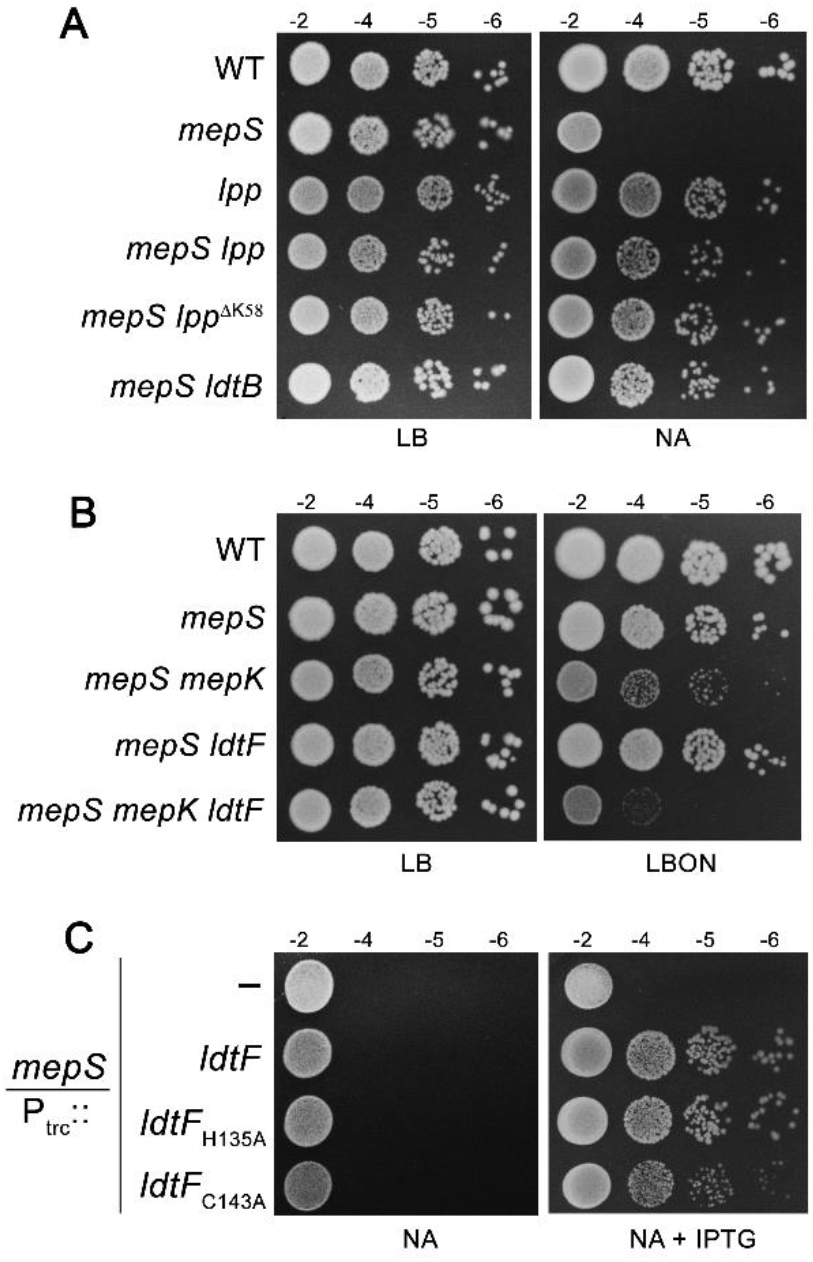
Genetic interactions of *mepS* with LDTs. (A) WT and its mutant derivatives carrying either deletion of *lpp*, *ldtB* or *lpp^ΔK58^*, an allele lacking C-terminal lysine were tested for viability at 37°C. (B) Indicated strains were grown and viability was tested as described above. (C) A *mepS* deletion mutant carrying either vector (pTrc99a; P_trc_::) or pRB1 (P_trc_::*ldtF*) or pRB2 (P_trc_::*ldtF*_H135A_) or pRB3 (P_trc_::*ldtF*_C143A_) were grown in LB broth supplemented with ampicillin and growth was examined on nutrient agar (NA) plates with or without IPTG (0.25 mM) at 37°C.

As these results intrigued us, we further investigated the role of LdtF by introducing the plasmids encoding each of the LDTs (26) into the *mepS* mutant. As shown in Fig. S3, plasmids encoding *ldtA*, -*B*, -*C*, -*D*, -*E* did not confer growth to *mepS* mutant whereas a plasmid encoding *ldtF* alone was able to moderately suppress the growth defects of *mepS* mutant. Another plasmid derivative carrying cloned *ldtF* downstream to an IPTG-dependent promoter (P_trc_::*ldtF*) also suppressed the growth defects of *mepS* mutant on NA (Fig. 2C) indicating that LdtF may have a distinct function compared to that of other LDTs. LdtF belongs to YkuD family of proteins and members of this family contain L,D-transpeptidase domain with an invariant cysteine residue at the active site (27). To further validate the role of LdtF, we constructed a mutant derivative with an alanine residue substituted for cysteine (LdtF-C143A) and examined its ability to suppress the *mepS* mutation. Fig. 2C shows that a plasmid encoding LdtF-C143A poorly suppressed the *mepS* deletion mutant whereas another variant coding for LdtF-H135A behaved like that of WT. However, deletion of *ldtF* alone in a WT strain did not confer any discernible phenotype when grown on LB, LBON (LB without NaCl) or NA plates at 30, 37 or 42°C except a slight reduction in the doubling time during exponential phase of growth (Fig. S4). In addition, *ldtF* deletion mutant did not exhibit significant sensitivity to any of the cell wall-antibiotics such as cephalexin, cefsulodin, mecillinam or vancomycin.

### LdtF modulates PG–Lpp linkages *in vivo*

To understand the basis of LdtF’s function, we examined the composition of PG in strains either having a deletion or multiple copies of *ldtF*. PG sacculi from these strains were prepared, digested with a muramidase (mutanolysin) followed by separation of soluble muropeptides by RP-HPLC and identification of the peaks by MS or MS-MS analysis (as described in Materials and Methods). No major difference was observed in the muropeptide profile of *ldtF* deletion derivative compared to that of WT (Fig. S5). In contrast, the PG sacculi of cells carrying additional copies of *ldtF* had considerable alterations (Fig. 3A; Table S3), the most significant being absence of peak 3 (tri-lys-arg) which is a disaccharide tripeptide attached to a lys-arg dipeptide (Fig. 3B, 3C). Tri-lys-arg muropeptides are generated due to the proteolytic activity of pronase which is used during preparation of PG sacculi to remove bound Lpp. Pronase cleaves Lpp at the 56^th^ position leaving the extreme C-terminal lys-arg dipeptide attached to the mDAP residue of the stem peptides resulting in generation of several species of muropeptides bound to lys-arg dipeptide (2). In addition to absence of peak 3, a muropeptide peak eluting at 46 min (labelled ‘Y’) was significantly elevated in the PG sacculi of cells carrying additional copies of LdtF (Fig. 3A). MS-MS analysis indicated this peak to be a tetra-tri dimer linked by 4–3 cross-bridge with a molecular mass of 1794 Da (Fig. 3B, 3C; Fig. S6). Absence of peak 3 with concomitant increase in peak 1 (a monomer of tri) allowed us to speculate that LdtF may have an ability to modulate the mDAP–Lpp linkages. Though the source of peak Y is not clear, it was not detected in a strain deleted for Lpp suggesting it may have originated by the activity of LdtF on PG–Lpp cross-links (Fig. S7A). Additionally, the incidence of peak Y was not dependent on the presence of functional LdtD and -E (Fig. S7B). Further, all other alterations observed due to overexpression of LdtF disappeared in a strain lacking Lpp reinforcing the suggestion that LdtF functions downstream of Lpp (Fig. S7A). As shown in Fig. 3A, analysis of PG in strains carrying plasmids encoding either LdtF-C143A or LdtF-H135A indicated a direct role for LdtF in modulation of mDAP–Lpp linkages (Table S3).

**Fig. 3.**
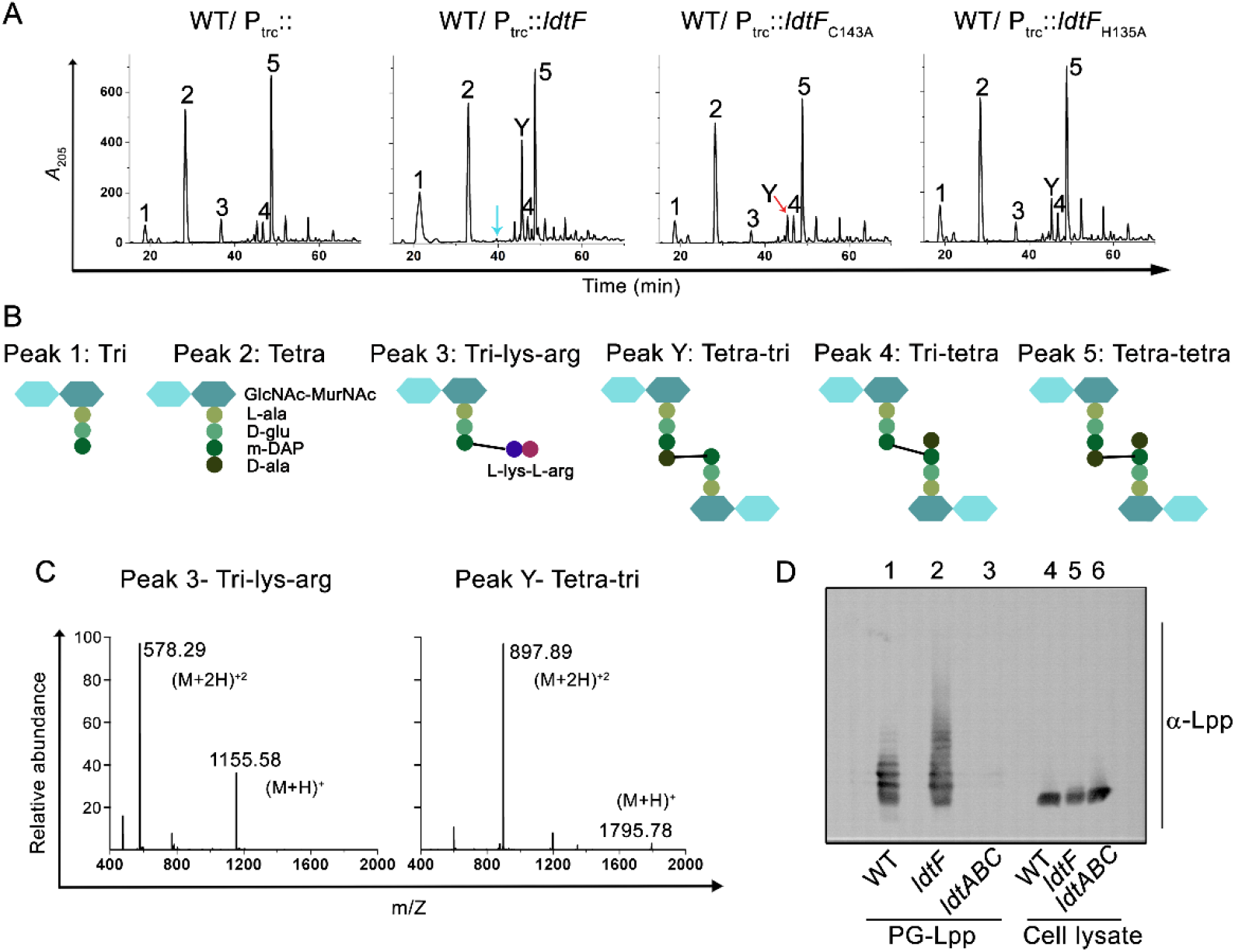
LdtF modulates PG–Lpp linkages. (A) HPLC chromatograms of PG sacculi of WT carrying either vector (P_trc_::), pRB1(P_trc_::*ldtF*), pRB2 (P_trc_::*ldtF*_H135A_) or pRB3 (P_trc_::*ldtF*_C143A_). Cultures were grown to an *A*_600_ of ∼1 in LB containing 0.2 mM IPTG followed by isolation and analysis of PG sacculi. (B) Structures of muropeptides (C) Mass spectra of peaks 3 and Y showing molecular mass (M+H)^+^ of 1,155.58 Da and 1795.78 Da (D) Determination of PG–Lpp linkages in WT, *ldtF* mutant was done by treating intact PG sacculi (with bound Lpp) with mutanolysin followed by electrophoresing the soluble muropeptides. Lpp containing muropeptides were visualized by western blot using anti-Lpp antibody. PG from *ldtABC* mutant was used as negative control. Cell lysates of WT, *ldtF* and *ldtABC* were used as controls (lanes 4, 5 and 6).

Because the above results implicated LdtF in regulation of PG–Lpp linkages, we examined the extent of these cross-links in cells lacking LdtF. To perform this experiment, Lpp-bound PG sacculi were prepared from WT and *ldtF* deletion mutant as described in Materials and Methods. PG sacculi from both strains were digested with mutanolysin, soluble muropeptides were collected and normalized amounts (Fig. S8) were electrophoresed using SDS-PAGE followed by western blotting and detection with anti-Lpp antibody. Fig. 3D shows that the PG sacculi derived from LdtF deletion mutant indeed contain a greater abundance of high molecular weight Lpp-bound muropeptides compared to that of WT, although the level of low-molecular weight Lpp-bound muropeptides were unchanged. This observation suggested an interesting possibility of LdtF specifically moderating the larger oligomeric Lpp–cross-linked muropeptides of the PG sacculus and the implications of this result are further discussed below.

### LdtF is a murein endopeptidase that cleaves PG–Lpp linkages

To examine the enzymatic activity of LdtF, a signal-less hexa-histidine tagged LdtF (LdtF^20-246^-His_6_) was overexpressed and purified as described in Materials and Methods. Treatment of soluble muropeptides derived from the PG sacculi of WT *E. coli* with purified LdtF yielded muropeptide fraction that totally lacked tri-lys-arg (peak 3) and tetra-tri-lys-arg (peak 6) with concomitant increase in tri- and tetra-tri muropeptides (Fig. 4A). Cleavage of tri-lys-arg and tetra-tri-lys-arg into tri- or tetra-tri muropeptides was also confirmed using purified fractions (Fig. 4B, 4C, 4D).

**Fig. 4.**
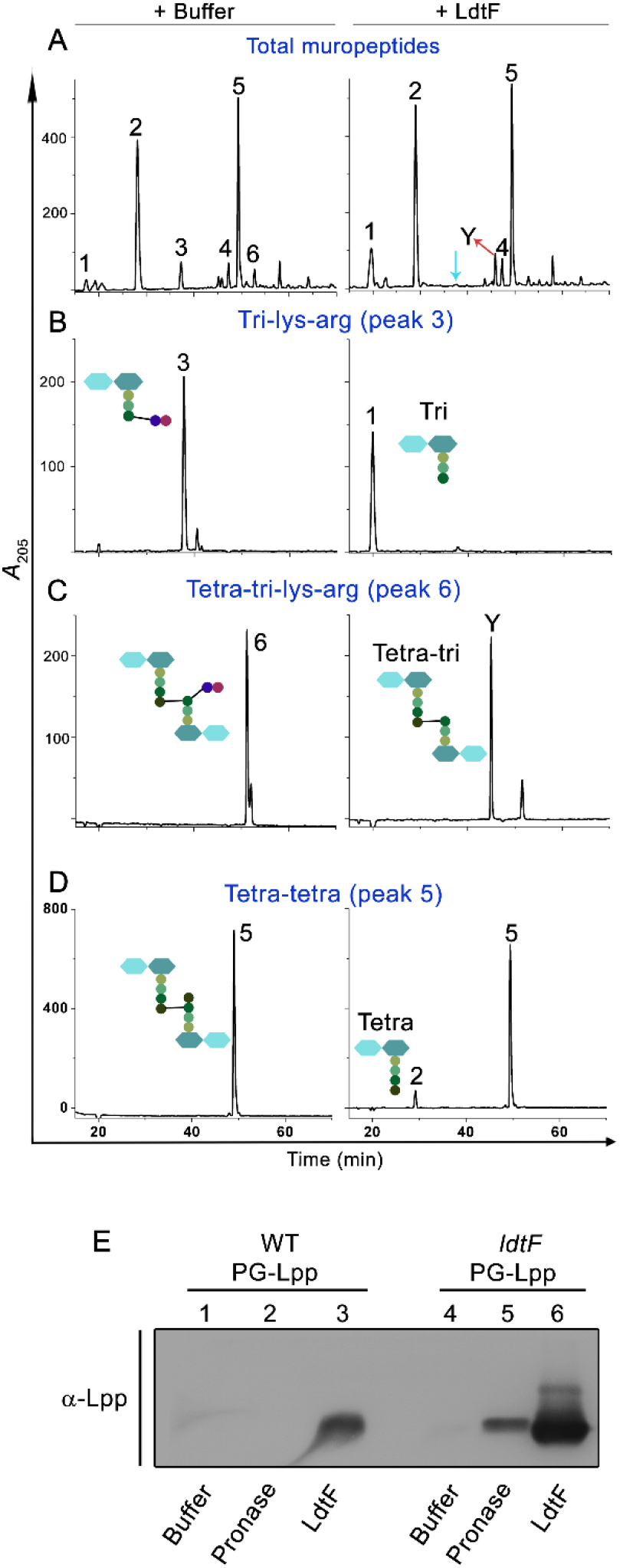
Endopeptidase activity of LdtF. (A) Soluble muropeptides of WT PG sacculi, (B) purified tri-lys-arg, (C) purified tetra-tri-lys-arg, or (D) purified tetra-tetra dimer were incubated either with buffer or LdtF (4 μM) for 16 h and separated by RP-HPLC. LdtF cleaved peak 3 (tri-lys-arg) to yield tri (peak 1); and peak 6 (tetra-tri-lys-arg) to yield tetra-tri (peak Y). LdtF showed an extremely weak activity on tetra-tetra dimer (peak 5) (E) Cleavage of Lpp protein from the PG sacculi (containing bound Lpp) of WT, *ldtF* mutant was tested by incubating the PG sacculi either with buffer, pronase (0.2 mg/ml) or LdtF (4 µM) for 16 h at 30°C. Pronase, a non-specific protease is used as a positive control.

We next examined the ability of LdtF to cleave the bound Lpp from the intact PG sacculi. To perform this experiment, Lpp-bound PG sacculi from WT and *ldtF* mutant were isolated and equal amounts of each were treated with purified LdtF. The soluble fraction was electrophoresed on SDS-PAGE and Lpp was detected by western analysis using anti-Lpp antibody. As a positive control, PG sacculi treated with pronase were used. Fig. 4E shows that both LdtF and pronase cleave Lpp from the PG sacculi and that the amount of Lpp released from the sacculi of LdtF deletion mutant was considerably higher (approximately 5-fold) than that of the WT (compare lanes 2 and 3 with 5 and 6). The remaining insoluble PG fraction was further analysed by RP-HPLC and as expected, the lys-arg muropeptides were not detected in LdtF-treated PG whereas pronase-treated PG contained the lys-arg muropeptides (Fig. S9). Overall, these results demonstrate the catalytic specificity of LdtF on PG–Lpp or PG–lys-arg substrates.

### LdtF is a glycine-specific carboxypeptidase that cleaves terminal glycine residue from the stem peptides

The above experiments clearly demonstrated hydrolytic activity of LdtF on PG–Lpp linkages formed by LdtA, -B and -C. LDTs also exchange the terminal D-ala of stem peptides with a glycine residue. To examine whether LdtF has any activity on glycine-substituted muropeptides, soluble muropeptides of a WT strain grown with glycine-supplementation were used as substrates for LdtF. As expected, growth of WT *E. coli* with exogenously added glycine resulted in accumulation of a large number of muropeptides with glycine at position 4 whose identity is determined by MS or MS-MS analysis (Fig. S10). Fig. 5A shows that LdtF effectively removes glycine from a variety of glycine-containing muropeptides. LdtF also cleaved several glycine-containing muropeptides prepared from PG sacculi of a strain overexpressing LdtD (Fig. 5B). Of these, three distinct types of glycine-containing muropeptides were purified to homogeneity and Fig. 5C, 5D, 5E show that LdtF removes glycine residue from all these muropeptides. However, the abundance of glycine-containing muropeptides remained the same in both WT and *ldtF* deletion mutant when grown with glycine-supplementation suggesting the existence of alternate carboxypeptidases that cleave the terminal glycine residue. In support of this idea, *ldtF* deletion mutant was not sensitive to addition of glycine and behaved just like that of WT strain.

**Fig. 5.**
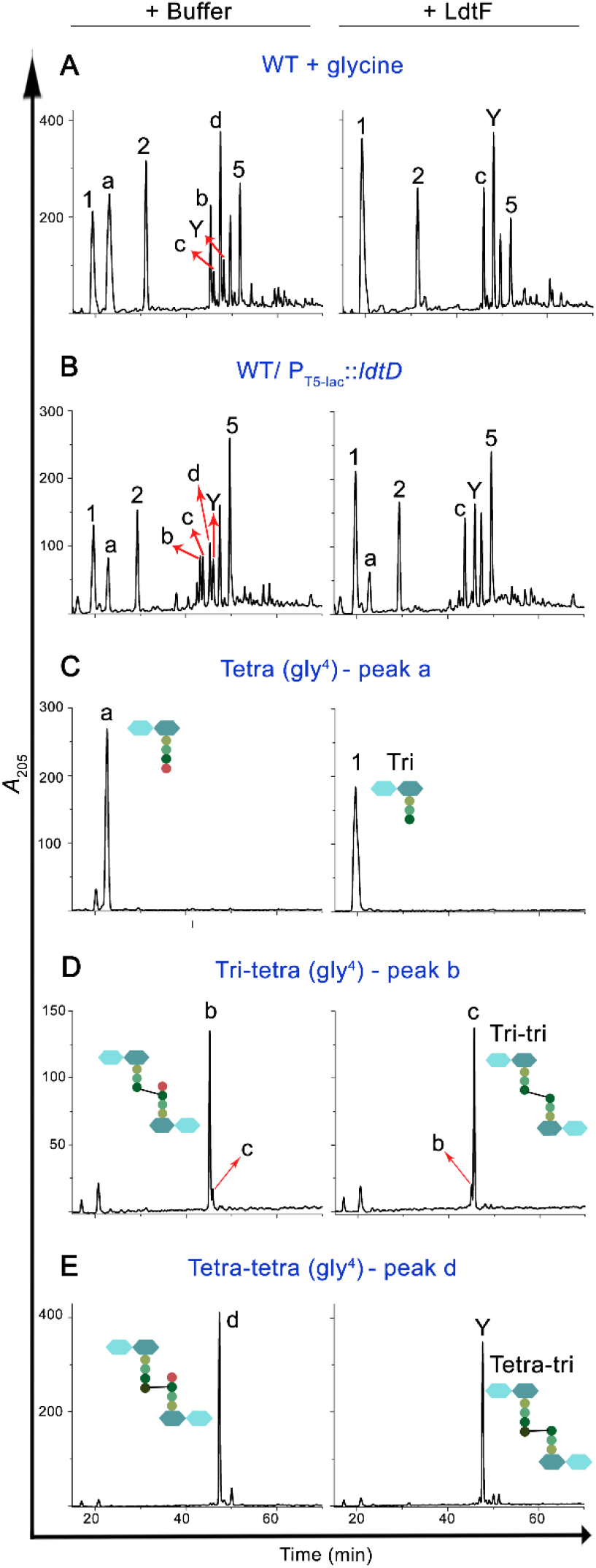
LdtF is a glycine-specific carboxypeptidase. (A) Soluble muropeptides generated from WT cells grown in minimal A medium (33) supplemented with 50 mM glycine, (B) soluble muropeptides of WT/ P_T5-lac_::*ldtD*, (C) purified tetra (gly^4^), (D) tri-tetra (gly^4^), or (E) tetra-tetra (gly^4^) were incubated either with buffer or LdtF (4 μM) and processed as described above. LdtF cleaved the terminal glycine residue completely from peak ‘a’ (tetra-gly^4^), ‘b’ (tri-tetra-gly^4^) or ‘d’ (tetra-tetra-gly^4^) to yield peak 1 (tri), ‘c’ (tri-tri) or Y (tetra-tri) respectively. All the muropeptides were analysed by mass spectrometry (Fig. S10).

Considering an earlier report that D,D-carboxypeptidases hydrolyse the terminal glycine from the stem peptides (2,28), we made a quadruple mutant deleted for major D,D-carboxypeptidases, DacA,-B, -C, -D and tested for sensitivity to glycine. As expected, the quadruple mutant formed smaller-sized colonies on glycine-supplemented media (Fig. S11). Introduction of *ldtF* deletion marginally exacerbated the defect of this quadruple mutant whereas multiple copies of *ldtF* moderately improved the growth of this mutant on glycine-containing media (Fig. S11) implicating a role for LdtF in removal of terminal glycine residue from the stem peptides. In sum, the above results demonstrate that LdtF is a glycine-specific carboxypeptidase.

### LdtF does not cleave NCDAAs from the stem peptides

To examine whether LdtF also cleaves the terminal NCDAA residues that are substituted by the exchange reaction of the LDTs, PG sacculi were made from WT *E. coli* grown in the presence of D-methionine, D-tryptophan or D-phenylalanine (22). Soluble muropeptides of these PG sacculi were separated and the peaks containing the NCDAA substitutions were identified by MS analysis (Fig. S12A) and used as substrates for LdtF (Fig. S12B). However, LdtF was not able to cleave the terminal NCDAA from any of these muropeptides (Fig. S12B).

### LdtF removes Lpp-mediated IM-PG linkages

Lpp is transported from the cytosol into the periplasm by Sec-mediated pathway and is eventually translocated to the OM by lipoprotein translocating machinery, LolABCDE (29). However, in certain transport-defective mutants, Lpp is stalled at the periplasmic face of the IM leading to the formation of IM-PG linkages by LDTs (30,31). It has been shown recently that absence of two small cytoplasmic membrane proteins, DcrB and YciB leads to mislocalization of Lpp at the IM, resulting in lethal IM-PG cross-links, and that this lethality is suppressed by deletion of either Lpp or LdtB (31). To examine the ability of LdtF to cleave the Lpp bound to IM, we made use of this mutant and observed that a deletion of *ldtF* exacerbates the growth defect of *yciB dcrB* double mutant (Fig. 6A). In addition, introduction of a multicopy *ldtF* plasmid (P_trc_::*ldtF*) partially restored the growth of the *yciB dcrB* double mutant, suggesting that LdtF may also cleave Lpp bound to the IM (Fig. 6B).

**Fig. 6.**
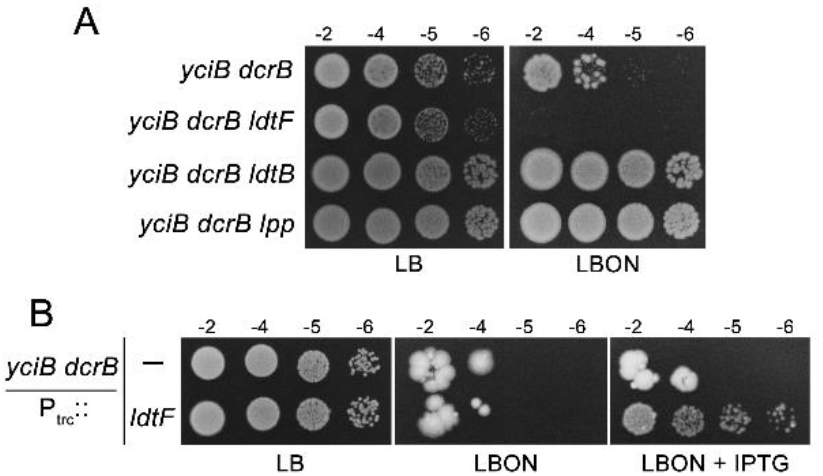
LdtF may also cleave Lpp-mediated IM-PG linkages. (A) *yciB dcrB* mutant and its *ldtF*, *ldtB* or *lpp* deletion derivatives were grown overnight in LB broth, serially diluted, and 5 µL of each dilution were spotted on indicated plates and tested for viability at 30°C. (B) Viability of *yciB dcrB* mutant carrying either vector (P_trc_::) or pRB1 (P_trc_::*ldtF*) was tested as described above at 37°C. IPTG was used at 0.1 mM.

## Discussion

Here, we report identification of a previously unknown peptidoglycan hydrolase, LdtF that cleaves Lpp (or Braun lipoprotein), an abundant OM lipoprotein which links OM to the PG sacculus in *E. coli*. LdtF is also a glycine-specific carboxypeptidase that removes the terminal glycine residue from the PG muropeptides. LdtF is a recently-identified member of YkuD family of proteins in *E. coli*; the other paralogs of this family comprise LdtA, -B, -C that catalyse the formation of mDAP–Lpp linkages and LdtD and -E which catalyse the formation of mDAP–mDAP cross-links. This study also represents an instance wherein members of a paralogous family perform contrasting but not overlapping functions.

### Role of LdtF in maintenance of envelope structure and stability

We identified LdtF in this study because its absence enhanced growth defects of a mutant lacking two of the PG elongation-specific endopeptidases, MepS and MepK and additionally, multiple copies of *ldtF* rescued the defects of *mepS* mutant (Fig. 2B, 2C). Moreover, we observed that absence of LdtF increases the PG–Lpp linkages (Fig. 3D) whereas more copies of *ldtF* decrease the level of PG-bound Lpp (Fig. 3A), suggesting a role for LdtF in modulating the degree of PG–Lpp cross-linkages. Subsequent biochemical analysis confirmed LdtF to be a hydrolase having two distinctive enzymatic functions- an L,D-endopeptidase activity that cleaves mDAP–Lpp cross-links and a carboxypeptidase activity that cleaves mDAP–gly linkages (Fig. 4; Fig. 5).

LdtF was earlier identified because a transposon insertion in *ldtF* (*yafK*) caused defective biofilm formation in an enteroaggregative *E. coli* strain (32). LdtF deletion has also been shown to confer additive sickness to a mutant defective in the transport of lipopolysaccharide (18). Although the basis of the above phenotypes is not clear, elevated OM-PG linkages may result in a defective cell envelope leading to such phenotypes. Excess OM-PG linkages may also alter the plasticity of the cell wall resulting in decreased fitness and survival of *E. coli*. Nevertheless, under laboratory conditions, absence of *ldtF* did not result in a large effect on the growth of *E. coli* excepting a small decrease in the doubling time (Fig. S4).

It is interesting to note that although the abundance of lower molecular weight Lpp-bound muropeptides was comparable in both WT and *ldtF* mutant (Fig. 3D, Fig. S5), the amount of bound Lpp is considerably higher in absence of LdtF (Fig. 3D, 4E). Occurrence of larger Lpp-bound oligomeric muropeptides in *ldtF* mutant (Fig. 3D) strongly suggests that LdtF may preferentially cleave PG–Lpp cross-links of higher order structures in the PG sacculus. Lpp-mediated OM-PG cross-linkages resulting in formation of large oligomers may distort the symmetry, or the organization of cell envelope and LdtF may perhaps work towards eliminating such linkages. In addition, the effect of *ldtF* alleles on the growth of mutants in which Lpp is stalled at the IM (Fig. 6) suggests a role for LdtF in removal of rare IM-PG linkages that may occur during transport of Lpp across the periplasm. LdtF may also facilitate PG turnover as the Lpp-linked muropeptides may not efficiently get recycled unless Lpp is cleaved from the PG sacculi.

Earlier studies implicated LdtF in facilitating the formation of 3–3 cross-links because ectopic expression of LdtF along with coexpression of LdtD or -E (in a strain lacking *ldtABCDEF*) increased the level of 3–3 cross-links (18–20). However, we did not observe increased 3–3 dimers when LdtF is overexpressed in WT; in contrast, we detected increased 4–3 linked dimer (peak Y) whose occurrence was dependent on Lpp but not on LdtD or-E (Fig. S7). The basis of this discrepancy is not clear – it may perhaps be due to the strain background used for these studies and needs to be further investigated.

### Regulation of PG–Lpp linkages

Lpp is an abundant OM lipoprotein (5,6,10) with one third of it covalently attached to PG, making almost 10% of the peptides linked to Lpp. However, it is not clear how *E. coli* maintains optimal levels of PG–Lpp linkages. The combined activities of LdtABC and LdtF may control the abundance of PG–Lpp linkages or alternatively structural/ conformational constraints of PG sacculi may limit the extent of PG–Lpp cross-link formation. These linkages are reported to be higher during conditions of stress including stationary-phase of growth and certain mutant conditions (2,16–18). LdtF promoter expression is shown to be higher in stationary-phase (18); however, preliminary experiments done in our laboratory to examine the endogenous LdtF expression (using a LdtF-FLAG fusion) show that the protein levels fall during stationary-phase suggesting a likely basis for the occurrence of higher amount of PG–Lpp linkages in stationary phase. It would be worthwhile to further examine how cells achieve a dynamic yet balanced level of PG-Lpp linkages to maintain the structural and functional integrity of the cell envelope.

### Role of PG–Lpp linkages in PG enlargement

Absence of PG–Lpp linkages either by deleting Lpp, LdtB or increasing the copy number of LdtF, partially rescued the growth defects of a *mepS* deletion mutant (Fig. 2A, 2C); however, the suppression was not very robust (Fig. S1, S2). In addition, other phenotypes of *mepS* such as sensitivity to β-lactam antibiotic, mecillinam or its synthetic lethality with deletion of *mepM* (23) were not suppressed by deletion of Lpp or LdtB suggesting that the absence of Lpp linkages may not significantly affect the process of PG enlargement. Lack of OM-PG tethering may perhaps alter the mechanical properties of the cell envelope and increase the flexibility of the PG sacculus, consequently resulting in a marginal growth advantage to *mepS* mutants in low osmolar conditions such as NA.

## Materials and Methods

### Media, bacterial strains and plasmids

LB has 0.5% yeast extract, 1% tryptone and 1% NaCl (33). LBON is LB without NaCl. NA (Nutrient Agar) has 0.5% peptone and 0.3% beef extract. Antibiotics were used at the following concentrations (µg/mL): ampicillin (Amp)-50, chloramphenicol (Cm)-30, and kanamycin (Kan)-50. Bacterial strains and plasmids used in this study are listed in Tables S1-S2 (SI).

### Molecular and genetic techniques

Recombinant DNA and plasmid constructions were performed as per standard methods. MG1655 genomic DNA was used as template and Phusion HF DNA polymerase was used for PCR amplifications and the plasmid clones were confirmed by sequence analysis. P1-phage mediated transductions and transformations were performed using standard methods (33). All strains are derivatives of MG1655 (Coli Genetic Stock Centre). Deletion mutations are from Keio mutant collection (34).

### Determination of PG–Lpp linkages in the PG sacculi

Lpp-bound PG sacculi were isolated from cultures of WT and Δ*ldtF* mutant, treated with mutanolysin and the soluble fraction was run on 15% SDS-PAGE followed by western blotting using anti-Lpp antibody.

### Determination of enzymatic activity

To examine the activity of LdtF, soluble muropeptides were incubated with either buffer or LdtF (4 µM) at 30°C for 16 h. Samples were heat inactivated, reduced with sodium borohydride and separated by RP-HPLC. Lpp cleavage was examined by incubating purified LdtF (4 µM) with the Lpp-bound PG sacculi for 16 h in 25 mM Tris-HCl (pH 8.0) at 30°C followed by electrophoresis of the soluble fraction on SDS-PAGE and western blotting with anti-Lpp antibody.

## Supplemental Information

Details of strains, plasmids, additional materials and methods are described in Supplementary Information (SI). Tables S1-S3 and Figures S1-S12 are included in SI.

## Acknowledgements

We thank NBRP: *E. coli* for Keio collection and ASKA plasmid library; Thomas Silhavy for Lpp-K58 allele and anti-Lpp antibody; Sujata Kumari for initiating the work; V Krishna Kumari and C Subbalakshmi for HPLC; B Raman and Y Kameshwari for mass spectrometry analysis; N Madhusudhana Rao for suggestions, and members of Reddy laboratory for critical comments. This work is supported by funds from Council of Scientific and Industrial Research (MLP0141) and Department of Biotechnology (Centre of Excellence in Microbial Biology), Govt of India (to MR). We acknowledge financial support from Department of Biotechnology (to RB) and University Grants Commission of India (to PKC).

## Author Contributions

RB and MR designed the study; RB and PKC performed the experiments; RB, PKC and MR analysed the data and wrote the manuscript.

## Supplemental Information

### Supporting Materials and Methods

#### Plasmid constructions

For PCR amplifications, genomic DNA of MG1655 strain was used as a template unless otherwise indicated. Amplification of DNA for cloning purposes was done using Phusion DNA polymerase (NEB) and clones obtained were confirmed by sequence analysis.

##### pRB1

The *ldtF* gene along with its native ribosome binding site (RBS) was PCR amplified using forward and reverse primers 5’-GC<u>TCTAGA</u>AGGAATAAGCAGTATGCGTAAA-3’ and 5’-CCC<u>AAGCTT</u>TTATTTTGCCTCGGGGAGCGTGT-3’ respectively and the resulting amplified DNA fragment was cloned using XbaI and HindIII sites (underlined in the primer sequence) in a cloning vector, pTrc99a to obtain pRB1. The clones were confirmed by sequence analysis and shown to suppress the NA-sensitivity of Δ*mepS* mutant at 37°C with 250 µM IPTG.

##### pRB2 and pRB3

To create site directed variants of *ldtF* (H135A and C143A), a 3 step PCR was performed. For this procedure, two primers were synthesized that are complementary to each other with desired mutations at the center (in bold and underlined). In the first PCR step, N-terminal fragment of *ldtF* gene was amplified using a common forward primer and a reverse primer containing the desired mismatch. In the second PCR step, C-terminal fragment of *ldtF* gene was amplified using a forward primer containing the desired mismatch and a common reverse primer. Desired mismatches code for an alanine instead of H135, and C143 in the LdtF. Common forward primer (containing its native RBS and an XbaI site), and reverse primer (containing a HindIII site) are used for cloning these variants.

Common forward primer: 5’-GC<u>TCTAGA</u>AGGAATAAGCAGTATGCGTAAA-‘3 and Reverse primer: 5’-CCC<u>AAGCTT</u>TTATTTTGCCTCGGGGAGCGTGT-3’.

The forward and reverse primers with nucleotide substitution are: Histidine to alanine change at codon 135- 5’-AAGGGAAATACCTGATGATC**<u>GCT</u>**GGCGATTGTGTTTCCATCGG-3’ and 5’-CCGATGGAAACACAATCGCC**<u>AGC</u>**GATCATCAGGTATTTCCCTT-3’

Cysteine to alanine change at codon 143- 5’-GCGATTGTGTTTCCATCGGC**<u>GCT</u>**TACGCAATGACCAATCAGGG-3’ and 5’- CCCTGATTGGTCATTGCGTA**<u>AGC</u>**GCCGATGGAAACACAATCGC-3’

In the third step, both the PCR products were mixed in 1:1 molar ratio and end filling was done by PCR in 10 cycles at low annealing temperature. After addition of common forward and reverse primer, PCR was resumed for the next 30 cycles. The final PCR product was digested with XbaI-HindIII and cloned into pTrc99a digested with the same enzymes. The recombinant plasmids, pRB2 (*ldtF*-H135A) and pRB3 (*ldtF*-C143A) were confirmed for the presence of mutations by sequencing.

##### pRB4

A fragment encoding LdtF^20-246^ was cloned into pET21b vector in between NdeI and XhoI sites using forward and reverse primers 5’-GGAATTC<u>CATATG</u>GGTTTGCTGGGCAGCAGTAG-3 and 5’-CCG<u>CTCGAG</u>TTTTGCCTCGGGGAGCGTGTAG-3’ respectively to generate a C-terminal 6XHis fusion vector. The plasmid was confirmed by sequencing and used for expression and purification of LdtF.

#### Construction of an *ldtF* deletion mutation (Δ149*ldtF*::Kan)

We constructed a partial deletion mutant of *ldtF* lacking N-terminal 1-149 amino acids using recombineering as described earlier (35). The hybrid primers used for constructing this deletion are:

FP: 5’- GTCCTGGCGTGTGTAACCGTTTTATCAAGGAATAAGCAGTATGTGTAGGCTGGAG CTGCTTC-3’

RP: 5’- CACCAGCGCACCAGTAACGAACTGGAATATCTCATCAATACCATATGAATATCCTCCTTAG-3’

In the first step, the Kan^R^ cassette of pKD4 vector (35) was amplified with above set of primers and the purified PCR product was electroporated into a strain encoding λ Red-Gam system. Transformants were selected on plates supplemented with 25 μg/ml kanamycin, followed by confirming the deletion by sequencing and linkage analysis. The deletion was subsequently transferred by P1 transduction into MG1655. This deletion mutant behaved exactly like that of the deletion mutant of Keio collection (in terms of causing sickness to *mepS mepK* double mutant and in PG composition) and hence, Keio deletion mutation was used throughout the study.

#### Construction of *ldtF*-FLAG fusion

Epitope tagging of LdtF using a 3xFlag was done at the 3′ end of the gene at its native chromosomal locus by recombineering as described earlier (36). The sequence of the hybrid primers used for construction of this fusion is given below:

FP: 5’- GCAGCCACAACTGGCATCAAACTACACGCTCCCCGAGGCAAAAGACTACAAAGACCATGACGGTG-3’ and,

RP: 5’- CGGGCAATGAAACCTGGCAAAAGATTATGCCAGGCGAATGGCGCCATATGAATATCCTCCTTAG-3’

The 3xFlag tag along with Kan^R^ cassette was amplified from plasmid pSUB11 (36) using the above primers. The 5’ end of (43 bases) forward primer has homology to the C-terminal of *ldtF* without stop a codon whereas the 3’ end (22 bases) has homology to the region encoding Flag epitope of pSUB11. Similarly, the 5’ end of reverse primer has a region homologous (42 bases) to the downstream sequence of *ldtF* whereas the 3’end has a region (22 bases) which has homology to the sequence of pSUB11 plasmid at the 3’ end. The PCR product was electroporated into DY378 and transformants were selected on plates supplemented with 25 μg/ml Kanamycin at 30°C on LB. The putative *ldtF-*3xFlag-Kan^R^ region was transferred into MG1655 by P1 transduction and the construct was confirmed by sequencing and linkage analysis. The expression of the fusion tag was confirmed by western blotting using anti-FLAG antibodies.

#### Confirmation of deletion mutation (mutant from Keio collection) of *ldtF*

The Keio deletion mutant of *ldtF* was confirmed by sequencing the gene-Kan^R^ junctions. The region encompassing the deletion mutation was amplified using the below primers and sequenced using the same primers.

FP: 5’- GACAGGCTTGCGTAAAACTC-3 and, RP: 5’- CAGGATGTGGAAATCGACTTCAGC-3

### Methods

#### Purification of LdtF

LdtF encoding plasmid, pET21b-*ldtF*^20-246^ (pRB4) was transformed into BL21 (λDE3) strain and transformants were selected on LB plates supplemented with ampicillin (Amp). One purified colony was grown overnight and used to dilute 1:100 into 50 ml fresh LB broth containing Amp. Culture was induced with 250 μM IPTG at 0.6 OD and further allowed to grow for 2 h at 37°C. Cells were harvested, and pellet was stored at –30°C. When required, pellet was resuspended in 1 ml of buffer (50 mM Tris-Cl, 300 mM NaCl and 20 mM imidazole, pH 8.0) and lysed by sonication. Cell debris was removed by centrifugation and the supernatant was mixed with 200 µl Ni^+2^-NTA agarose (Qiagen) and mixed for 1 h at 4°C. This mixture was loaded onto pre-washed empty column (Bio-Rad) and washed with 30 ml wash buffer-1 (50 mM Tris, 300 NaCl, 30 mM imidazole, pH 8.0) and 20 ml of wash buffer-2 (50 mM Tris, 300 NaCl, 50 mM imidazole, pH 8.0). The bound proteins were eluted with 5 ml of elution buffer (50 mM Tris, 300 NaCl, 150 mM imidazole, pH 8.0) and concentrated to 2.5 ml using a 3 kDa cut-off centrifugal membrane filter (Millipore). This was then loaded onto a buffer exchange PD-10 column and protein was eluted into 3.5 ml 2 x storage buffer (100 mM Tris, 200 mM NaCl, 2 mM DTT). The fraction was further concentrated to 250 μl using 3 kDa cut-off centrifugal membrane filter and mixed with equal volume of 100% glycerol and stored at –30°C.

##### Viability assays and Microscopy

Viability of the indicated strains was examined by growing cultures overnight, serially diluting (10^−2^, 10^−4^, 10^−5^ and 10^−6^), and placing 3-5 µl aliquots of each dilution onto the required plates. Plates were normally incubated for 18-24 h at the indicated temperature. To measure growth rates, overnight grown cultures were diluted 1:100 into fresh medium, allowed to grow and OD at 600 nm was determined after every 1 h interval and data were plotted using Origin software. For microscopy, immobilized cultures on a thin 1% agarose pad were visualized using Zeiss AxioImager.Z2 microscope by DIC.

##### Preparation of PG sacculi

Isolation of PG was done as described earlier (2,24). Cells grown to *A*_600_ of 1.0 were collected by centrifugation at 10,000×g for 10 min at 4°C. Cell pellet (from 1000 ml) was resuspended in 6 ml of ice-cold deionized water and added drop wise into 6 ml of boiling 8% SDS with vigorous stirring followed by boiling for another 45 min to solubilize membranes and to destroy high molecular weight DNA. After overnight incubation at room temperature, the PG sacculi were collected by high speed centrifugation (200,000×g, 40 min) and washed thoroughly with deionized water to completely remove SDS. High molecular weight glycogen and covalently bound lipoprotein, Lpp were removed by treating with α-amylase (100 µg/ml in 10 mM Tris-HCl, pH 7.0, 2 h at 37°C) and pre-digested pronase (200 µg/ml, 90 min at 60°C). Enzymes were inactivated by boiling with equal volume of 8% SDS for 15 min and pure sacculi were obtained by ultracentrifugation and washed several times with water till SDS was completely removed. Final pellet was resuspended in 0.5 ml of 25 mM Tris-HCl (pH 8.0) and stored at –30°C. Lpp-bound PG sacculi were prepared from cultures grown up to *A*_600_ of ∼6.0 using the above-described protocol except that the treatment with Pronase was not done.

##### Analysis of PG sacculi

PG analysis was done as previously described (2,24). Essentially, the sacculi were digested with 10 U mutanolysin (Sigma-Aldrich) at 37°C in 25 mM Tris-HCl (pH 8.0) for 16 h and soluble muropeptides were collected after centrifugation. This fraction was reduced with 1 mg of sodium borohydride in 50 mM sodium borate buffer (pH 9.0) for 30 min and excess borohydride was destroyed by addition of 20% phosphoric acid. pH was adjusted to 3–4 and samples were loaded onto a reverse phase C18 column (Zorbax 300 SB; 250×4.6 mm, 5 mm) connected to Agilent technologies RRLC 1200 system. Column temperature was 55°C and binding was done at a flow rate of 0.5 ml/min with 1% acetonitrile in water containing 0.1% trifluoroacetic acid (TFA) for 10 min. Muropeptides were eluted in a gradient of 1–10% acetonitrile containing 0.1% TFA at a flow rate of 0.5 ml/min for next 60 min (using RRLC online software called Chemstation). Absorbance of muropeptides was detected at 205 nm.

##### Mass spectrometry (MS) analysis of muropeptides

Muropeptide fractions collected during HPLC were dried and reconstituted into 5% acetonitrile with 0.1% formic acid and loaded onto a reverse phase PepMap^TM^ RSLC - C18 column (3 µm, 100Å, 75µmx, 15cm) connected to Q-Exactive^TM^ HF Hybrid Quadrupole-Orbitrap^TM^ Mass Spectrometer (Thermo Fisher Scientific, USA). Peaks were analysed by tandem MS and structures were decoded based on molecular mass of the individual fragments.

#### Determination of Lpp-bound muropeptides

Normalized muropeptides were boiled with laemmli loading dye and separated using 15% SDS-PAGE. Lpp was detected by western blot using rabbit anti-Lpp antibody (rabbit). As a positive control (free form of Lpp) cell lysates of WT, Δ*ldtF*, or Δ*ldtABC* (equivalent of 0.012 OD) were used.

#### Cleavage of Lpp from intact PG sacculi

Lpp-bound PG sacculi were isolated from WT and Δ*ldtF* (grown upto ∼6 OD) without pronase treatment as described. Equal volumes of the PG sacculi were treated with buffer, pronase (as positive control) or LdtF (4 µM) for 16 h at 30°C. Reaction was stopped by heat inactivation and supernatant was collected after centrifugation at 15000xg for 15 min. These fractions were run on 15% SDS-PAGE to detect the Lpp released from intact PG sacculi. The remaining pellet fraction was further digested with mutanolysin and the resulting soluble muropeptides were separated by RP-HPLC.

### Supplementary Figures

**Fig. S1.**
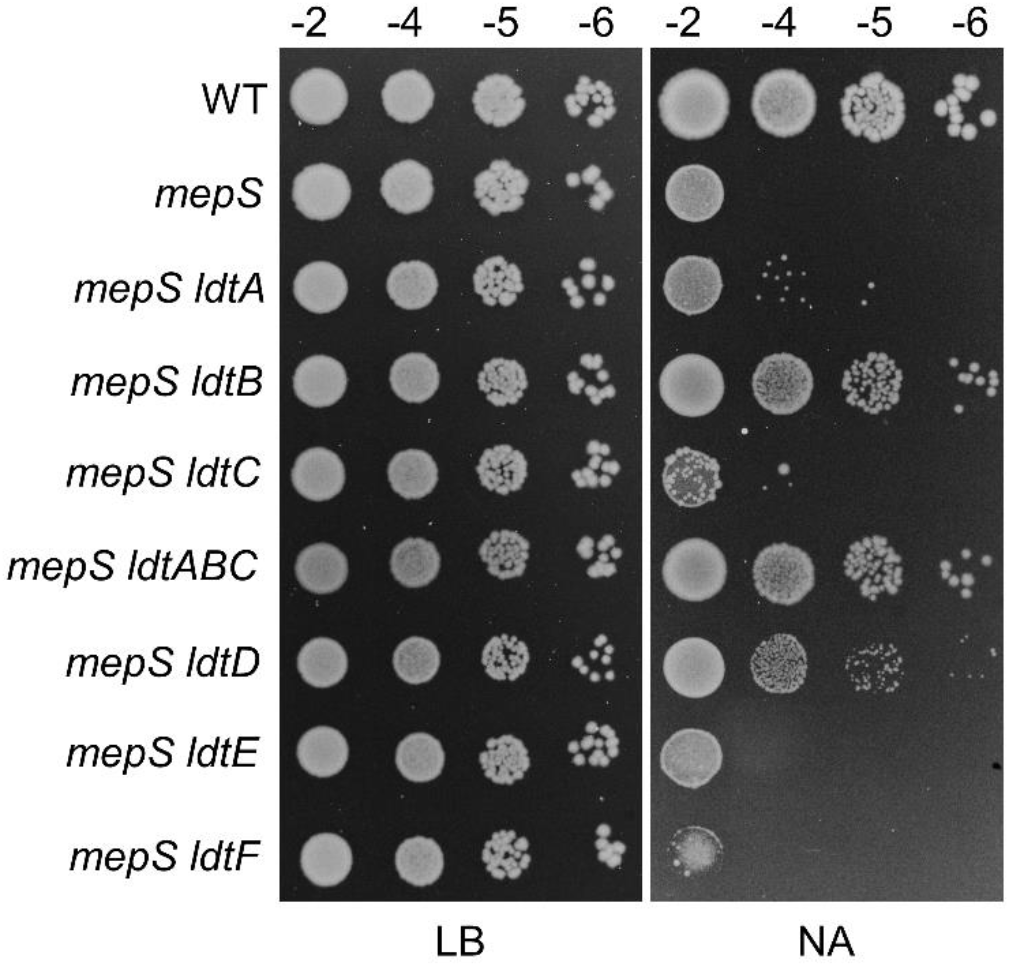
Genetic interactions of *mepS* with *ldts*. WT (MG1655) and its deletion mutant derivatives were grown overnight in LB broth, serially diluted, and 5 µL of each dilution were spotted on indicated plates and tested for viability at 37°C.

**Fig. S2.**
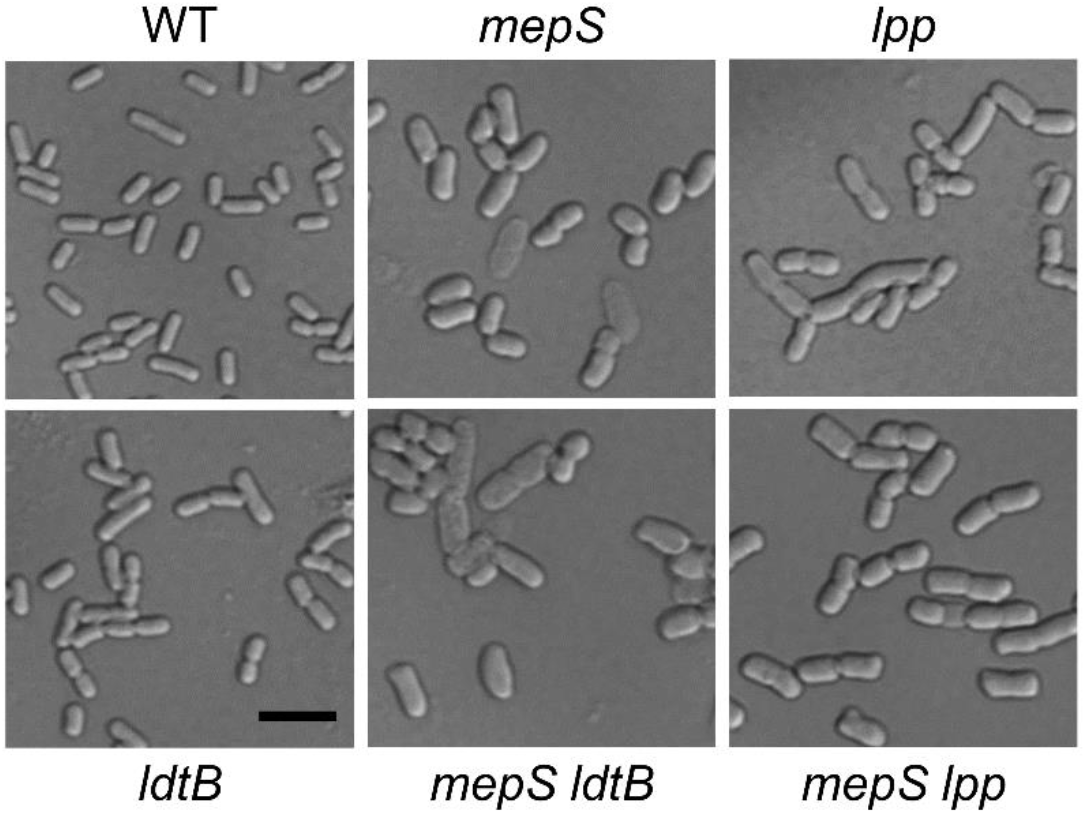
Microscopic images of WT and its various mutant derivatives. Indicated strains were grown overnight and diluted 1:500 into prewarmed Nutrient Broth and grown till *A*_600_ of 1.0 at 37°C and visualized with DIC microscopy. The scale bar represents 5 μm.

**Fig. S3.**
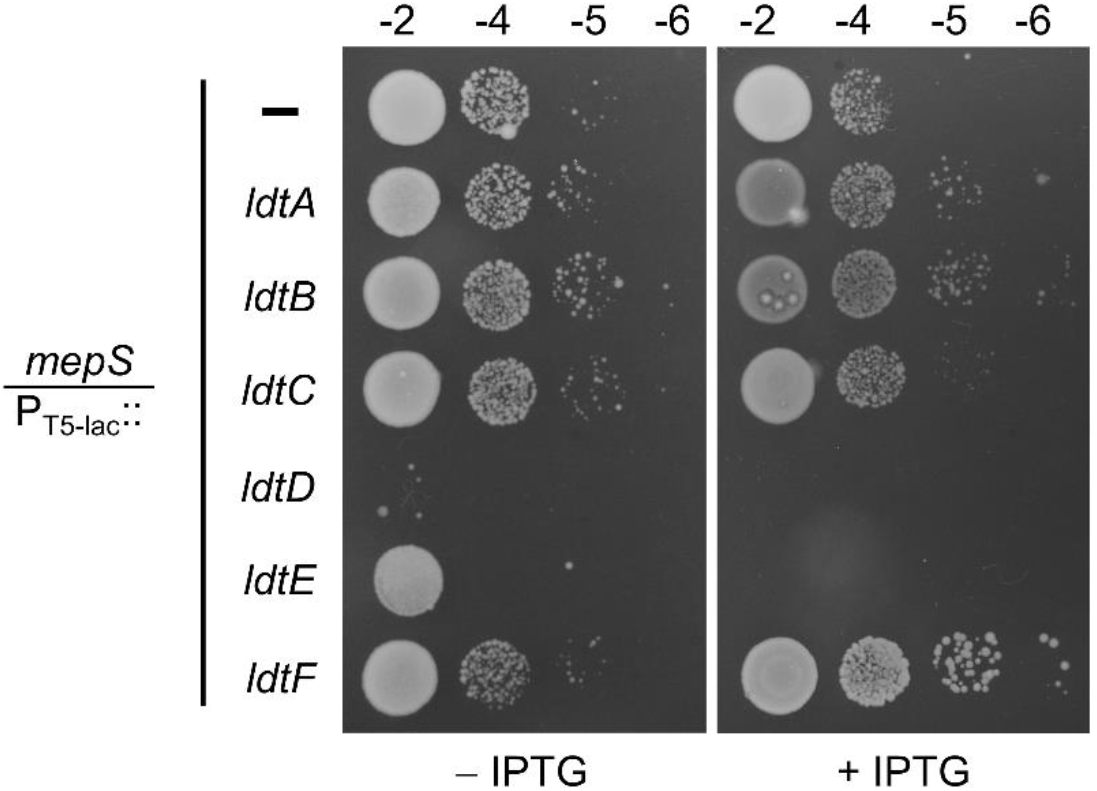
Effect of multiple copies of Ldts on growth of *mepS* mutant. Δ*mepS* mutant carrying either pCA24N vector (P_T5-lac_::) or its derivatives (P_T5-lac_::*ldtA-F*) were grown and viability was tested on NA plates at 37°C with or without 50 µM IPTG.

**Fig. S4.**
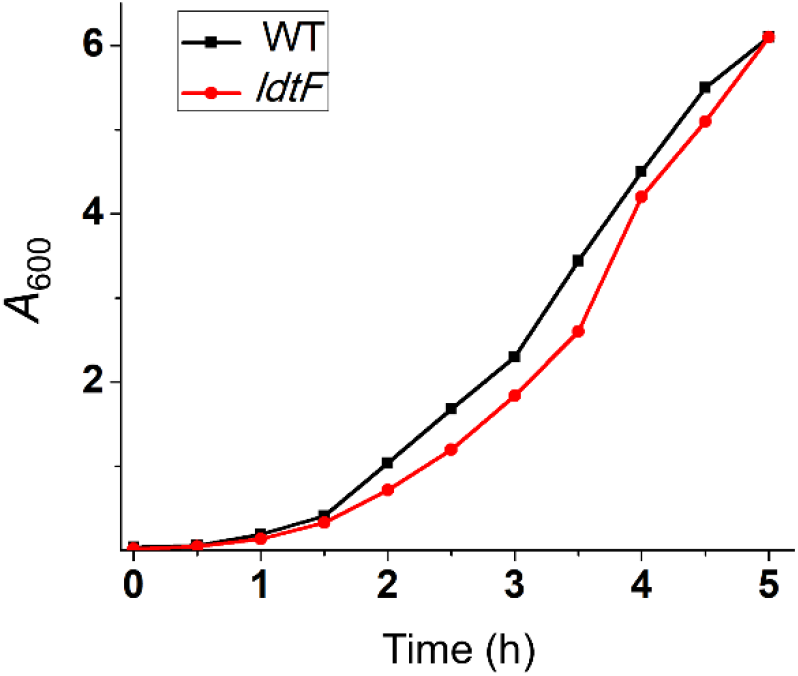
Growth curve of *ldtF* mutant. Overnight grown cultures of WT and *ldtF* were sub-cultured 1:500 into fresh LB and growth was monitored every 30 min at 37°C. In addition, *ldtF* mutant shows comparable decrease in the doubling time compared to that of WT when grown in any media including LBON or Minimal A media with or without glycine-supplementation.

**Fig. S5.**
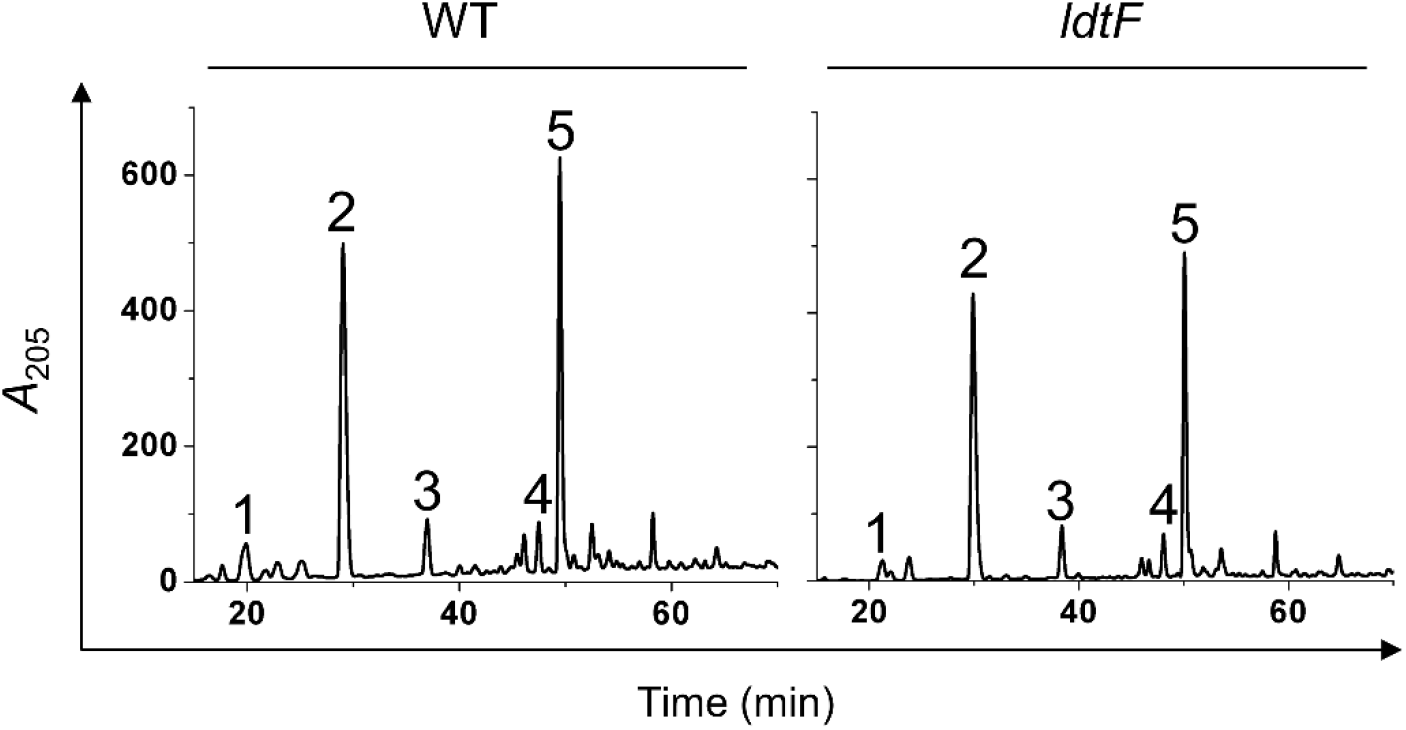
PG composition of *ldtF* mutant. HPLC chromatograms of PG sacculi isolated from WT and *ldtF* mutant. Strains were grown to an *A*_600_ of ∼1 in LB followed by isolation and analysis of PG sacculi. Data shown is representative of three independent experiments.

**Fig. S6.**
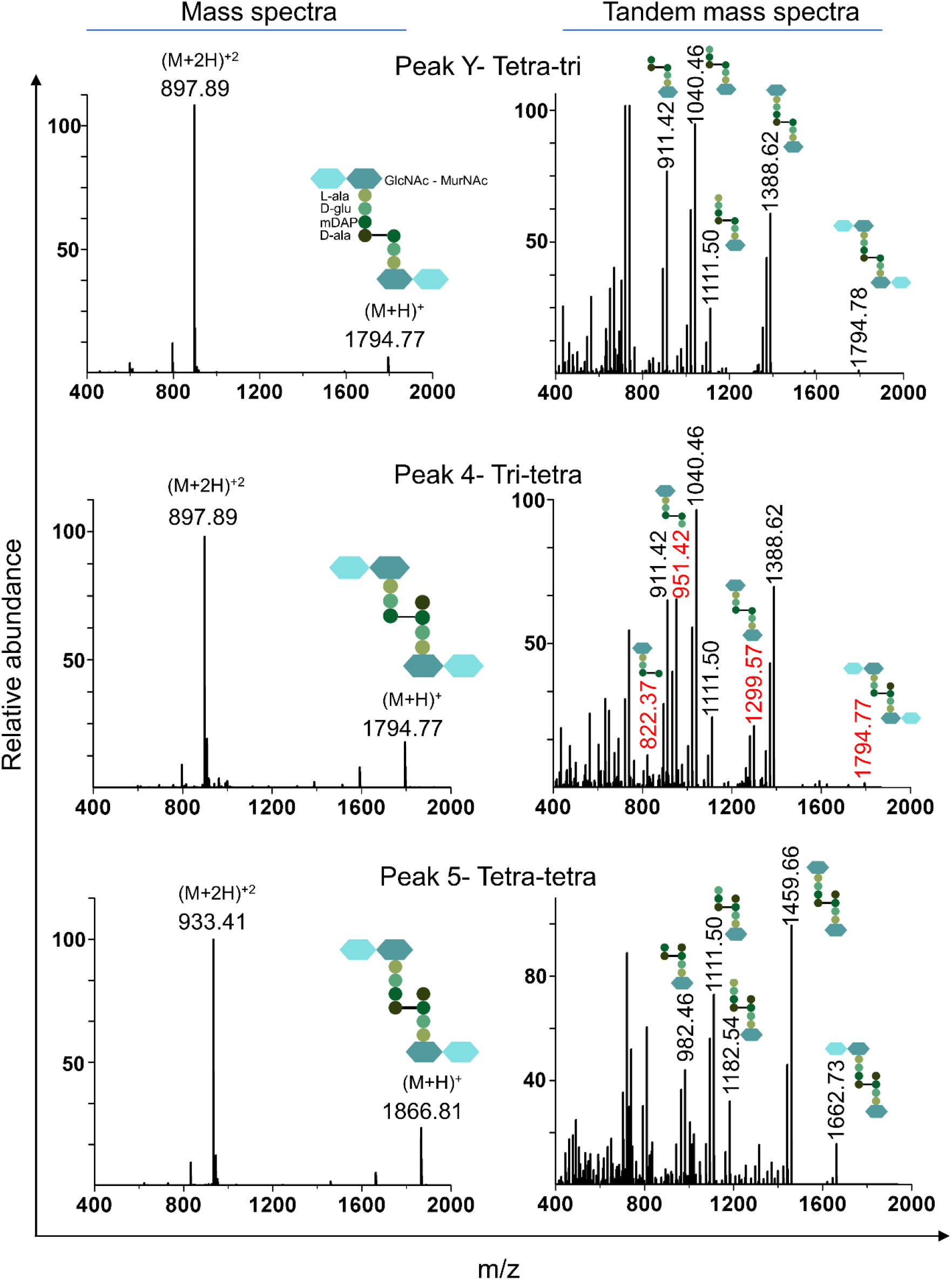
Mass spectrometry analysis of various muropeptides including tetra-tri, tri-tetra and tetra-tetra. Mass spectra of various muropeptides indicating the molecular mass (both +1 and +2 charges) is shown. The signature peaks indicating the 3–3 cross-links in tri-tetra are highlighted in red.

**Fig. S7.**
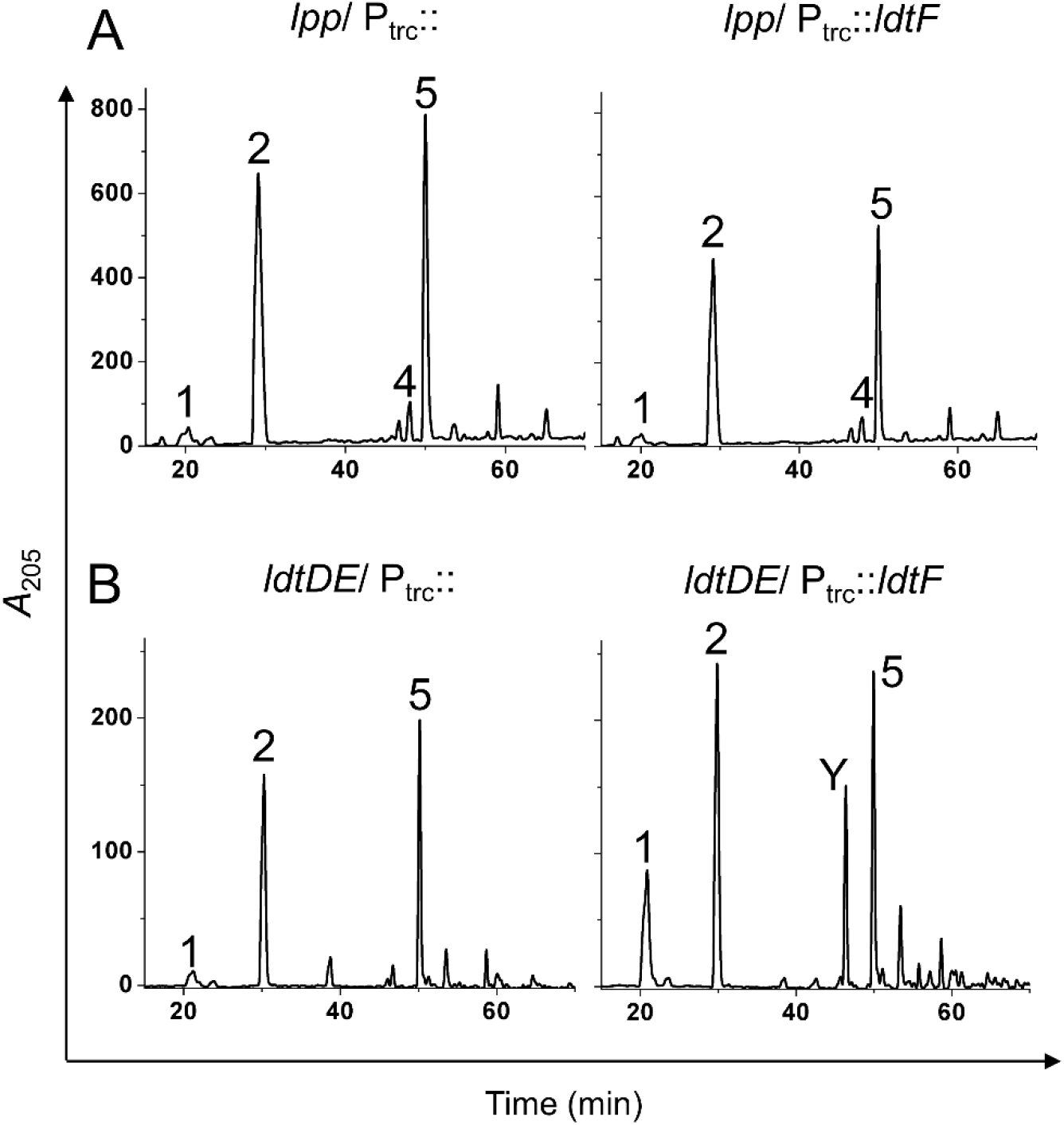
PG composition of strains with ectopic expression of LdtF. HPLC chromatograms of PG sacculi isolated from (A) *lpp* or (B) *ldtDE* deletion mutants carrying either vector (pTrc99a; P_trc_::) or pRB1(P_trc_::*ldtF*). Strains were grown to an *A*_600_ of ∼1 in LB containing 150 µM IPTG followed by isolation and analysis of PG sacculi. Note that in absence of Lpp, increased LdtF does not alter the PG composition. On the other hand, the effect of LdtF is independent of LdtD and -E.

**Fig. S8.**
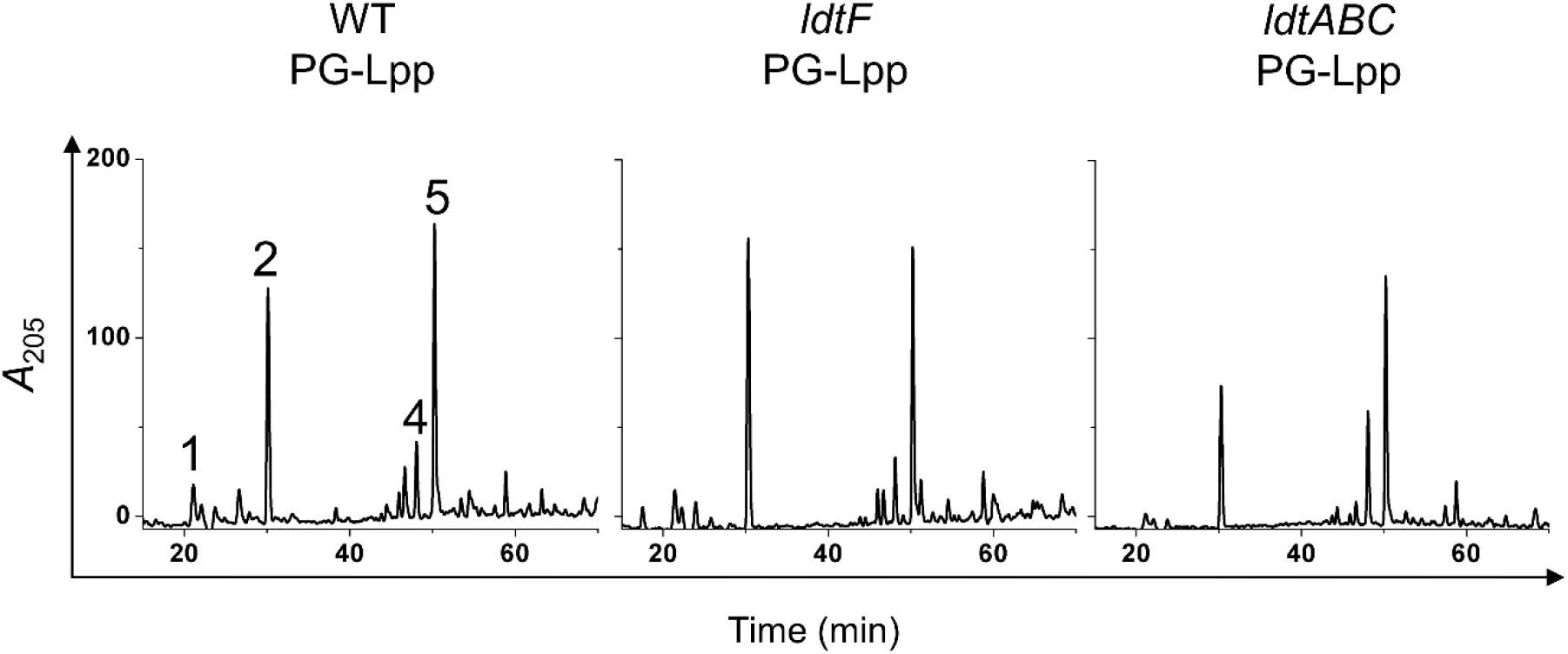
HPLC chromatograms of Lpp-bound PG sacculi from WT, *ldtF* and *ldtABC* mutant. The data obtained from these chromatograms was used to normalize the amount of sample loaded for performing the experiment described in Fig. 3D.

**Fig. S9.**
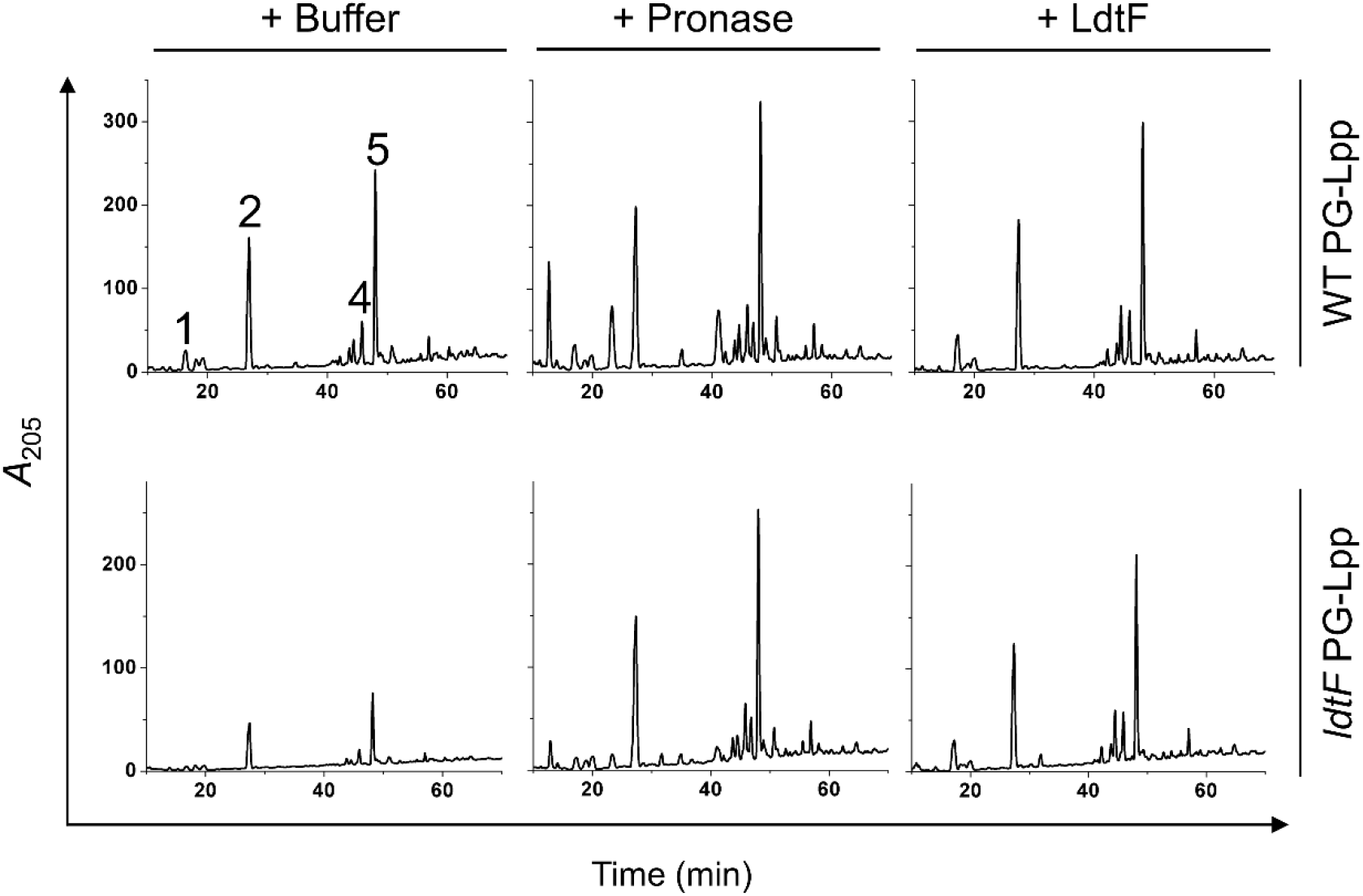
HPLC chromatograms of Lpp-bound PG sacculi from WT and *ldtF* mutant. The PG sacculi were treated either with buffer, pronase (0.2 mg/ml) or LdtF (4 μM) for 16 h at 30°C and the remaining insoluble fraction was treated with mutanolysin and separated by HPLC. Note that the tri-lys-arg peak is absent in LdtF-treated PG sacculi.

**Fig. S10.**
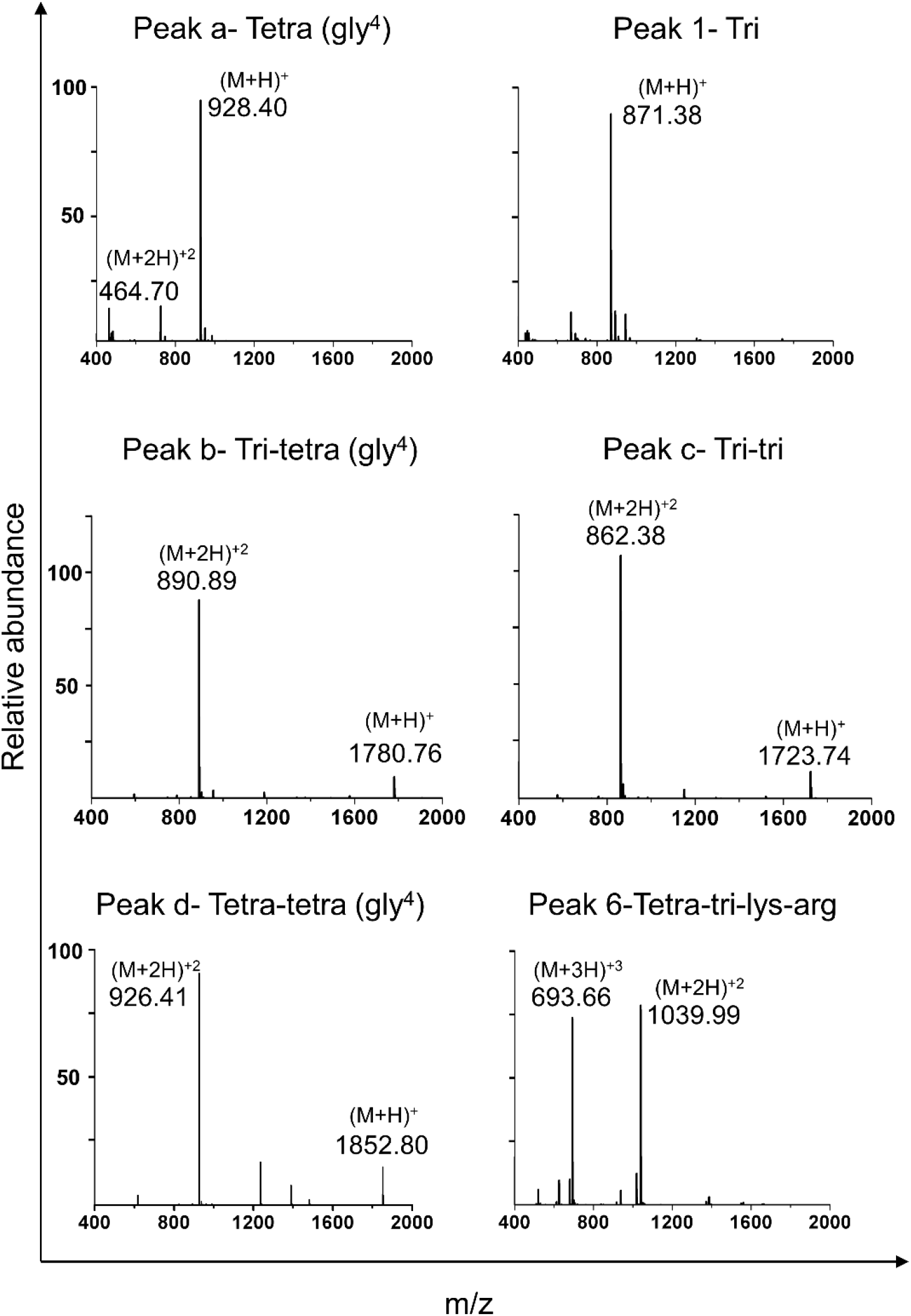
Identification of muropeptides using Mass spectrometry analysis. Mass spectra of various muropeptides indicating the molecular mass (both +1 and +2 charges) is shown.

**Fig. S11.**
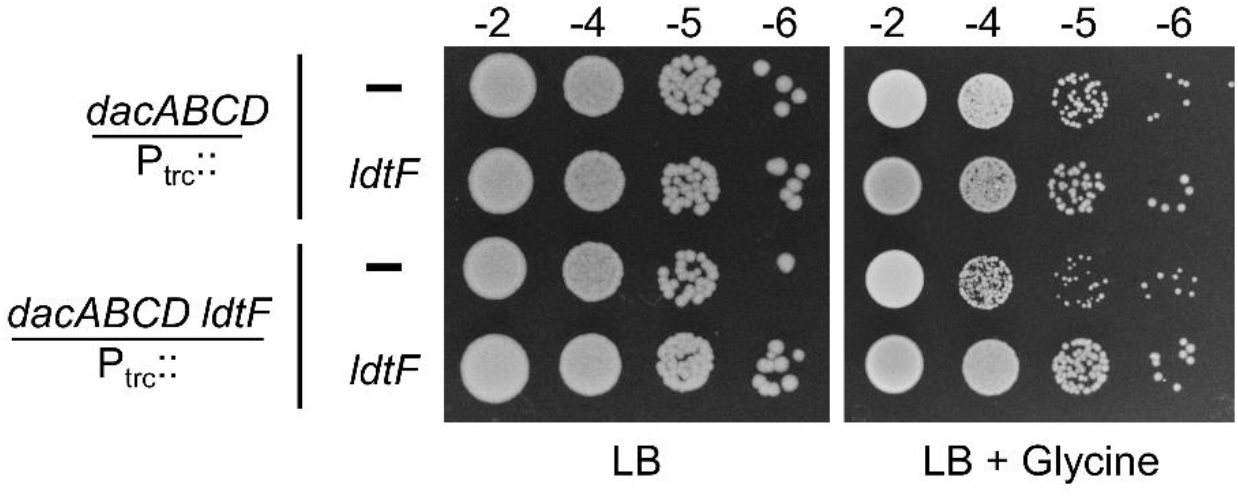
Effect of LdtF on glycine-toxicity of *dacABCD* mutant. Cultures of *dacABCD* and *dacABCD ldtF* mutants carrying either vector (pTrc99a; P_trc_::) or pRB1(P_trc_::*ldtF*) were grown in LB broth, serially diluted, and 5 µL of each dilution were spotted on indicated plates and tested for viability at 37°C. Glycine and IPTG are used at 25 mM and 100 µM respectively. *dacABCD* deletion mutant was slow-growing on glycine-supplementation and *ldtF* deletion exacerbated the sickness whereas more copies of *ldtF* marginally improved the growth of the quadruple mutant.

**Fig. S12.**
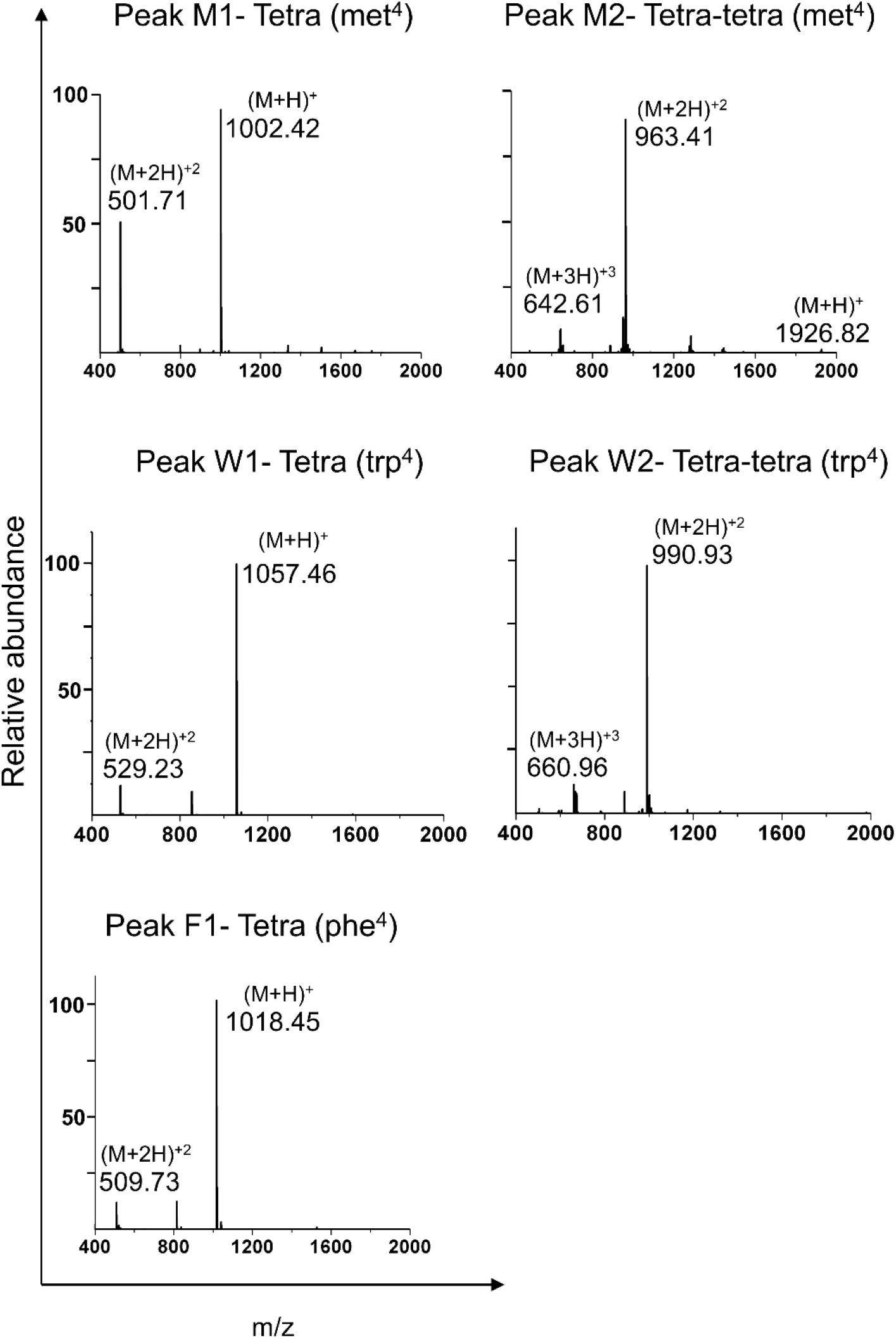
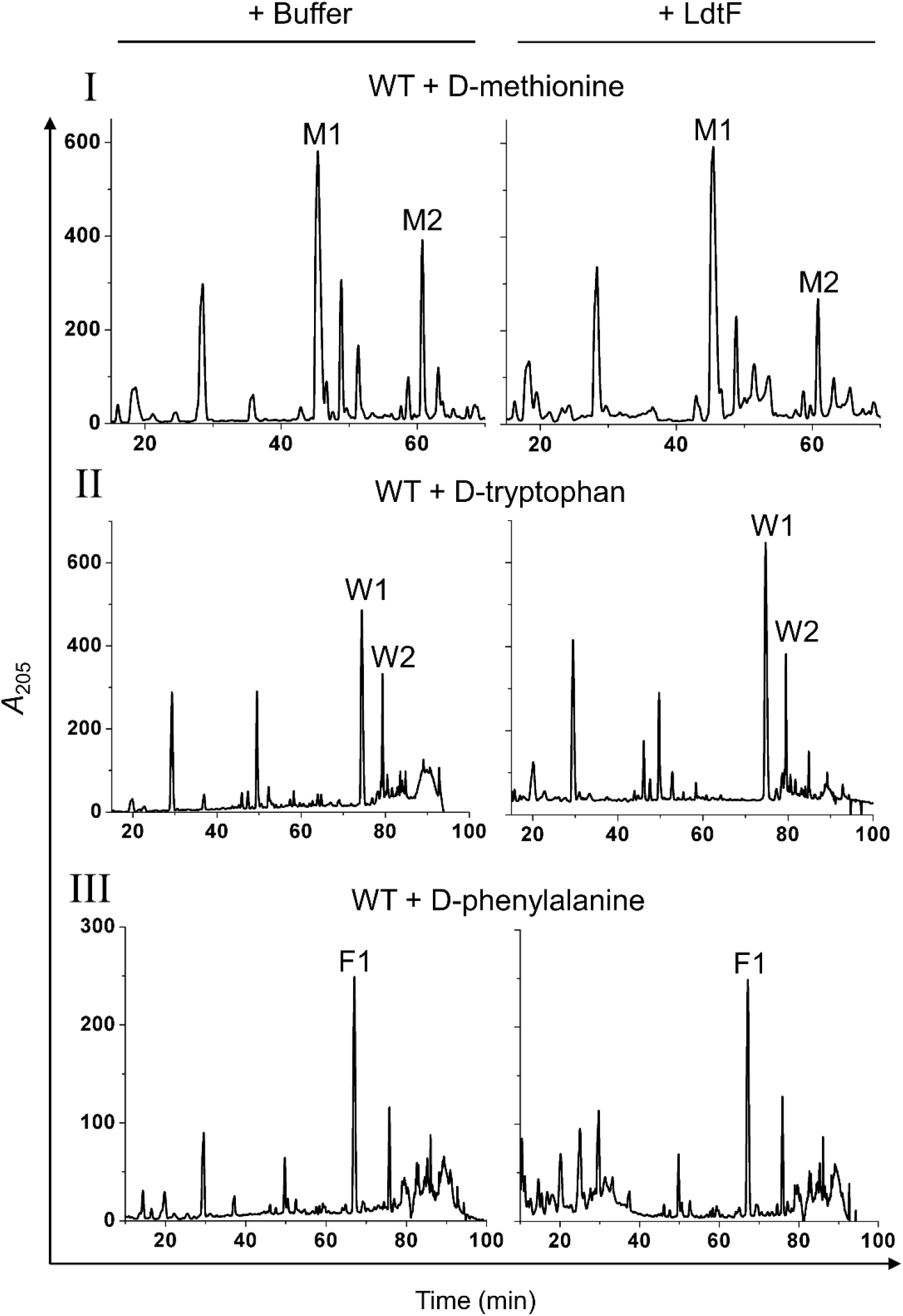
Effect of LdtF on NCDAA containing muropeptides. (A) Mass spectrometry analysis of muropeptides containing NCDAA (D-methionine, D-tryptophan or D-phenylalanine) (B) Treatment of soluble muropeptides from PG sacculi of strains grown in 20 mM D-methionine (I), 15 mM D-tryptophan (II), or 15 mM D-phenylalanine (III) were incubated either with buffer or LdtF (4 μM) for 16 h and separated by RP-HPLC. LdtF did not cleave the terminal D-amino acids from any of the NCDAA-muropeptides.

**Table S1.**
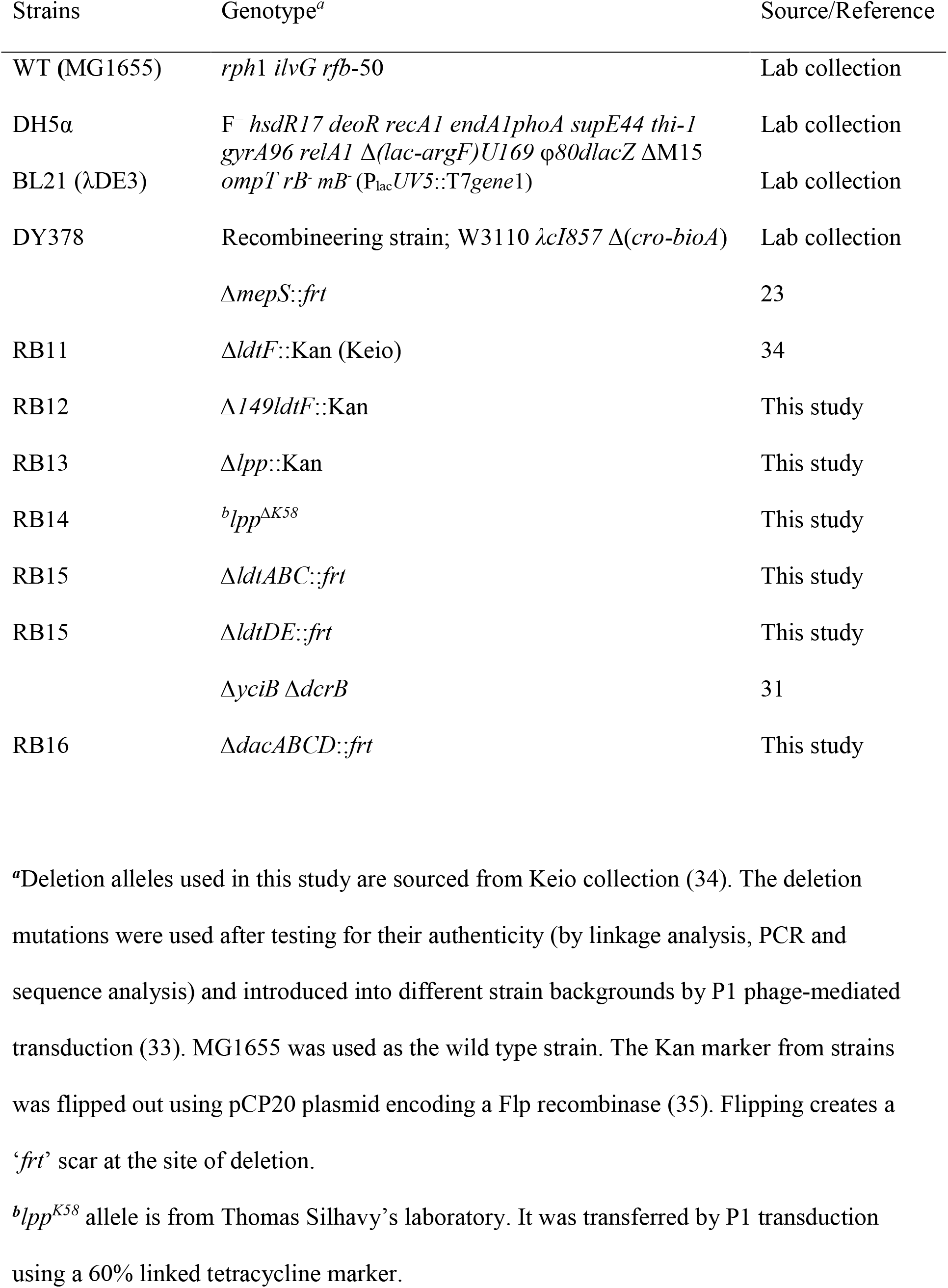
Strains used in this study.

**Table S2.** Plasmids used in this study.

| Plasmid | Relevant features | Source/Reference |
| --- | --- | --- |
| pET21b | ColE1, Amp<sup>R</sup>, *lacIq*, T7*lac* | Lab collection |
| pTrc99a | ColE1, Amp<sup>R</sup>, *lacIq*, P<sub>trc</sub> | Lab collection |
| pCA24N | Cm<sup>R</sup>, *lacI<sup>q</sup>*, P<sub>T5-lac</sub> | 26 |
| pCA24N-*ldtA* | Cm<sup>R</sup>, *lacI<sup>q</sup>*, P<sub>T5-lac</sub>::*ldtA* | 26 |
| pCA24N-*ldtB* | Cm<sup>R</sup>, *lacI<sup>q</sup>*, P<sub>T5-lac</sub>::*ldtB* | 26 |
| pCA24N-*ldtC* | Cm<sup>R</sup>, *lacI<sup>q</sup>*, P<sub>T5-lac</sub>::*ldtC* | 26 |
| pCA24N-*ldtD* | Cm<sup>R</sup>, *lacI<sup>q</sup>*, P<sub>T5-lac</sub>::*ldtD* | 26 |
| pCA24N-*ldtE* | Cm<sup>R</sup>, *lacI<sup>q</sup>*, P<sub>T5-lac</sub>::*ldtE* | 26 |
| pCA24N-*ldtF* | Cm<sup>R</sup>, *lacI<sup>q</sup>*, P<sub>T5-lac</sub>::*ldtF* | 26 |
| pRB1 | pTrc99a-*ldtF* | This study |
| pRB2 | pTrc99a-*ldtF*-H135A | This study |
| pRB3 | pTrc99a-*ldtF*-C143A | This study |
| pRB4 | pET21b-*ldtF*<sup>20-246</sup> | This study |

**Table S3.** Muropeptide composition of strains carrying multiple copies of *ldtF*.

| Muropeptide<br>(Peak) | % Area of muropeptide peaks<sup>a</sup> |  |  |  |
| --- | --- | --- | --- | --- |
|  | WT/<br>P<sub>trc</sub>:: | WT/<br>P<sub>trc</sub>::*ldtF* | WT/<br>P<sub>trc</sub>::*ldtF*<sub>H135A</sub> | WT/<br>P<sub>trc</sub>::*ldtF*<sub>C143A</sub> |
| Tri (1) | 5.81 ± 0.06 | 15.74 ± 0.47 | 9.51 ± 0.47 | 8.91 ± 1.41 |
| Tetra (2) | 38.91 ± 0.37 | 34.19 ± 1.70 | 35.14 ± 1.42 | 37.6 ± 1.47 |
| Tri-lys-arg (3) | **4.57 ± 0.53** | **0.7 ± 0.60** | **2.2 ± 1.22** | **1.86 ± 1.06** |
| Tetra-tri (Y) | **3.0 ± 0.23** | **11.75 ± 2.67** | **5.34 ± 0.06** | **5.27 ± 0.77** |
| Tri-tetra (4) | 3.54 ± 0.35 | 2.47 ± 0.45 | 3.99 ± 0.25 | 4.87 ± 0.18 |
| Tetra-tetra (5) | 32.22 ± 0.70 | 25.15 ± 1.79 | 27.44 ± 2.6 | 26.33 ± 2.91 |

<sup>a</sup>Muropeptide analysis was done by calculating the relative percentage area of each muropeptide from the HPLC chromatograms.

